# Organoid-to-organoid transplantation and pro-neurogenic treatment promote graft integration and neuronal repair in a human ischemic model

**DOI:** 10.64898/2026.09.25.754353

**Authors:** Ricardo Gómez-Oliva, Sergio Helgueta, Nerea Alonso-Galeron, Alessia Cambier, Fatemeh Ghanavatinejad, Gerald Masset, Lisa Collin, Pedro Nunez-Abades, Rosario Hernández-Galán, Carmen Castro, Ira Espuny-Camacho

## Abstract

Neuronal loss resulting from brain injury and neurodegenerative diseases currently lacks effective regenerative therapies. While neurogenic compounds and cell transplantation hold promise, clinical translation is hindered by poor graft integration and limited cell survival. Here, we developed a novel organoid-to-organoid transplantation model using human pluripotent stem cell-derived cortical organoids (hCOs) to evaluate a dual cell transplantation and pro-neurogenic strategy for brain repair. We demonstrate that pro-neurogenic treatment with ER272 stimulates early-stage hCO growth and enhances neurogenesis through a PKC- and Notch-dependent pathway. Mechanistically, ER272 upregulates genes associated with extracellular matrix, cell adhesion and growth factor signaling. Notably, combining early-stage hCO transplantation with ER272 treatment significantly enhanced graft integration and improved neuronal repair in an ischemic-like model of late-stage host hCOs.

This novel human organoid-to-organoid platform offers a robust translational tool to test regenerative therapies and highlight the potential of combining cell transplantation with pro-neurogenic stimulation for the treatment of neurological conditions associated with neuronal loss.

## INTRODUCTION

Brain injury caused by neurodegenerative disease, trauma or ischemia frequently results in neuronal loss and persistent cognitive impairment. Several strategies have been developed to promote brain repair, including the delivery of growth factors to stimulate endogenous neurogenesis, the reprogramming of non-neuronal cells into neurons and the transplantation of embryonic tissue or stem cell-derived neural progenitors ^1–9^.

Strategies focusing on pharmacological stimulation of endogenous neurogenesis are being explored clinically for psychiatric disorders, neurodegenerative diseases and traumatic brain injury. However, their clinical efficacy has so far been modest ^3–7,10^, partly because of differences in dosing and insufficient or variable delivery across the blood-brain barrier and/or transient or limited effects ^7,11,12^. This limitation suggests that complementary strategies capable of replacing lost cells may be required to repair the damaged human brain. Pro-neurogenic diterpenes have recently emerged as a potentially attractive alternative, as they stimulate neural stem cell (NSC) proliferation both *in vitro* and *in vivo* and can reach the brain following intranasal administration in mouse models ^13–16^. One example is ER272, a plant-derived 12-deoxyphorbol ester diterpene that activates members of the protein kinase C (PKC) family through binding to their regulatory domains, thereby mimicking the physiological PKC activator diacylglycerol (DAG) ^17,18^. Its proliferative activity relies on epidermal growth factor receptor (EGFR) signaling and involves phosphorylation and metalloprotease-dependent cleavage of EGFR ligands, followed by their release to the extracellular environment ^19–22^. Previous studies showed that ER272 promotes proliferation and neurogenesis in the adult mouse brain under both physiological^14^, and aging ^15^ conditions through the activation of PKCs and the release of the transforming growth factor α (TGFα) and to a lesser extent neuregulins. Intranasal administration of ER272 also improved cognitive performance in senescence-accelerated mouse-prone 8 (SAMP8) mice, supporting its ability to reach the brain and induce functional neuronal benefits *in vivo*. However, whether ER272 elicits comparable responses in the human brain remains unknown.

Alternative approaches such as cell transplantation rely on a direct exogenous supply of neural cells to repair the damaged brain. Clinical transplantation trials initiated in the late 1980s and 1990s used human brain embryonic tissue to treat neuronal loss in Parkinson’s disease patients. Whereas these studies provided proof-of-principle evidence that transplantation of embryonic tissue can produce functional benefits, they also highlighted substantial limitations, including restricted tissue availability, ethical concerns, variable graft composition and limited procedural standardization ^23–25^. Pluripotent stem cell-derived neural progenies may provide a more reproducible and scalable alternative. We and others have shown that cortical progenitors derived from human embryonic stem cells (hESCs) or human induced pluripotent stem cells (hiPSCs) can integrate into the injured adult mouse brain, differentiate into cortical neurons, receive host innervation and repair cortical axonal projections and synaptic connections with proximal and distal cortical targets ^1,2,26–29^, showing their potential as brain repair strategies in preclinical animal models. Ongoing clinical trials are currently evaluating hESC- or hiPSC-derived neuronal cell products for the treatment of neurological conditions, including Parkinson’s disease, ischemic stroke and traumatic brain injury. However, translation from animal models to patients remains challenging. Variability in donor-cell differentiation, limited graft survival and integration, and the inhibitory environment of the injured adult human brain may restrict therapeutic efficacy ^5,30^. These limitations suggest that successful brain repair may require not only an appropriate source of donor cells, but also pharmacological modulation of the host and graft environments to support donor-cell expansion, maturation and integration.

Advances in human pluripotent stem cell differentiation have enabled the generation of three-dimensional brain models, including human cortical organoids (hCOs), that reproduce key cellular and molecular features of human corticogenesis *in vitro* ^31–34^. Cortical organoids have been used to study developmental brain pathologies such as microcephaly ^31^, autism spectrum disorders (ASD) ^35^ and Timothy syndrome ^36^, as well as the neurological consequences of ZIKA or SARS-CoV2 infection, among others ^37–39^. They have also been adapted to model hypoxic and ischemic injury through oxygen and/or glucose deprivation (OGD) ^40–42^. These models offer a human-relevant platform to investigate the efficacy of regenerative drug treatment for the diseased brain. However, to date, there are no human *in vitro* models that recapitulate cell transplantation and pro-neurogenic compound treatment for brain repair.

Here, we used hESC- and hiPSC-derived hCOs to test the effects of ER272 pro-neurogenic treatment on human cortical cells. We found that whereas ER272 treatment leads to increased proliferation and neurogenesis rate in early-stage hCOs, this effect is largely absent in late-stage hCOs exposed to ischemic-like injury. To overcome this limited endogenous response, we developed a novel *in vitro* organoid-to-organoid transplantation model to test a dual strategy to regenerate an ischemic-like injury in a human cortical model. Combined cell transplantation and ER272 treatment led to improved graft integration and recovery of lost cortical neurons and neuronal fibers within the injured host organoid. Together, our findings indicate that effective repair of injured human cortical organoids requires both the provision of new neural cells and the pro-neurogenic modulation of the host environment.

## RESULTS

### Diterpene ER272 promotes the growth and proliferation of early-stage human cortical organoids

The diterpene ER272 has been shown to increase neural cell proliferation in neurogenic niches in mouse models *in vivo* ^14,15^, however, its proliferative effect in human cells remains unknown. To analyze the effect of the diterpene ER272 in human brain cells, we exposed hESC-derived early-stage hCOs to ER272 for 1 week starting at day 14 (Fig. 1A). ER272 increased total organoid area (Fig. S1A-B) and the proportion of neural progenitors SOX2^+^, proliferative Ki67^+^ and mitotic PHH3^+^ cells when compared to vehicle-treated hCOs (Fig. 1B-E). Moreover, ER272-treated hCOs showed a higher proportion of S-phase EdU^+^ cells than controls, consistent with increased cell proliferation (Fig. 1B,F). In agreement with this data, we found an increase in the number of ventricular zone-like (VZ-like) neuronal progenitor regions and in VZ-like area (Fig. 1G-I) together with higher Nestin and SOX2 protein levels in ER272-treated hCOs compared to controls (Fig. 1J-K).

**Figure 1.**
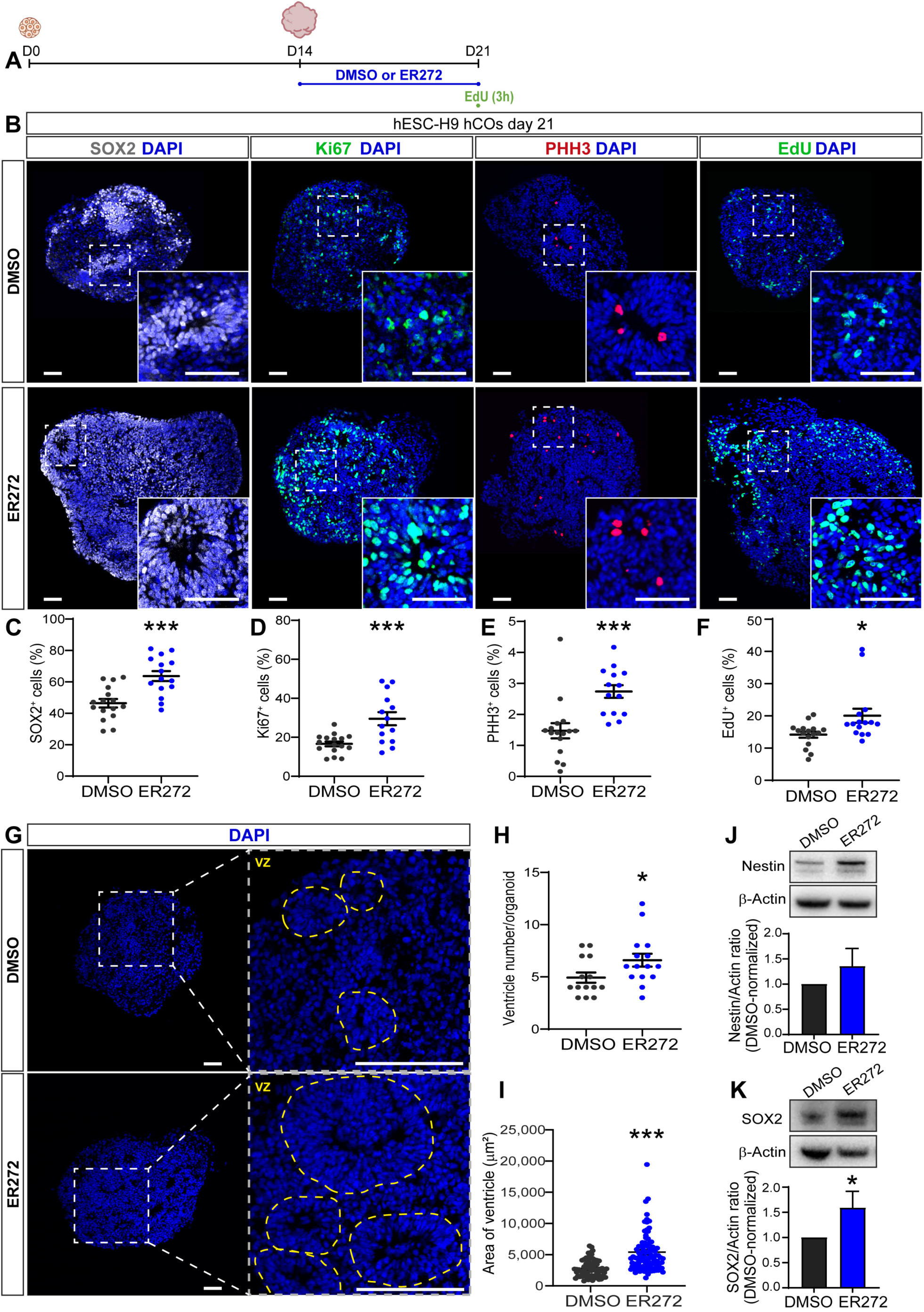
Diterpene ER272 promotes the growth and proliferation of early-stage human cortical organoids. **(A)** Schematic representation of the generation of H9 hESC-derived early-stage hCOs, treated with DMSO (vehicle) or ER272 from day 14 (D14) to day 21 (D21), followed by a 3 h EdU pulse before collection at D21. **(B)** Images and high magnification confocal views of immunostained H9 hESC-derived hCOs at day 21 showing the neural progenitor marker SOX2 (grey), the proliferative marker Ki67 (green), the mitotic marker PHH3 (red) and EdU^+^ (green) cells. Nuclei were counterstained with DAPI (blue). **(C-F)** Quantification of the percentage of SOX2-, Ki67-, PHH3- or EdU-positive cells relative to the total number of DAPI^+^ cells in DMSO- or ER272-treated H9 hESC-derived early hCOs. Data are presented as mean ± SEM from three independent experiments (n=14-16 organoids per condition). **(G)** Images of ventricular zone-like (VZ) regions identified by DAPI staining in DMSO- or ER272-treated H9 hESC-derived hCOs. **(H)** Number of VZ-like regions per organoid from three independent experiments (n=14-15 organoids per condition). **(I)** Area of VZ-like regions from three independent experiments (n=68-99 ventricles). **(J, K)** Western blot and quantification of Nestin and SOX2 protein levels in DMSO- or ER272-treated H9 hESC-derived hCOs at day 21. Protein levels were normalized to β-Actin and expressed relative to the DMSO control. Data are shown as mean ± SEM from one experiment (n=3 organoid samples per condition). *p < 0.05 and ***p < 0.001, Student’s t test. Scale bars, 100 μm.

Together, these data show that ER272 expands the neural progenitor pool and promotes the growth of early-stage hCOs.

### ER272 treatment increases neurogenesis in early-stage human cortical organoids

To determine whether ER272 treatment ultimately increases neurogenesis in hCOs, as previously observed in mouse models ^14,15,43^, we examined different pan-neuronal and cortical layer-specific markers at the end of the treatment (day 21; Fig. 2A left), or 1-week after treatment withdrawal (day 28; Fig. 2A right). This analysis revealed an increase in the expression of general neuronal markers such as MAP2 and TUBB3 (Fig. 2B-C,E-F,I-J and Fig. S1C-D), and increased proportion of CTIP2^+^ layer V and TBR1^+^ layer VI cortical cells (Fig. 2B,D-E,G and Fig. S1C,E) in ER272-treated hCOs compared to control hCOs. Next, we compared the rate of cell cycle exit between ER272-treated and control hCOs after a 24h EdU-pulse followed by removal of EdU and culture for 10 days (EdU chase; Fig. 2A right). These experiments showed an increase in total number of EdU^+^ cells in ER272-treated hCOs at day 28 (Fig. S1F-G), implying a higher number of post-mitotic cells 1-week post-treatment. Remarkably, we observed a higher percentage of CTIP2^+^ EdU^+^ cells within the EdU^+^ population in ER272-treated cells (Fig. 2E,H), demonstrating a higher percentage of layer V cortical neuronal production following ER272 treatment. In addition, we sought to determine if ER272 pro-proliferative effects were maintained post-treatment following the removal of diterpens from the hCO culture media. We found that hCOs analyzed 7 days after ER272 withdrawal had no differences in the number of proliferative Ki67^+^ and late G2/M-phase PHH3^+^ cells when compared to control hCOs (Fig. 2K-M), suggesting that the proliferative effect of ER272 did not persist after treatment withdrawal. These findings were also reproduced in two independent hiPSC-derived hCO lines (GM38 and GM40) (Fig. S2).

**Figure 2.**
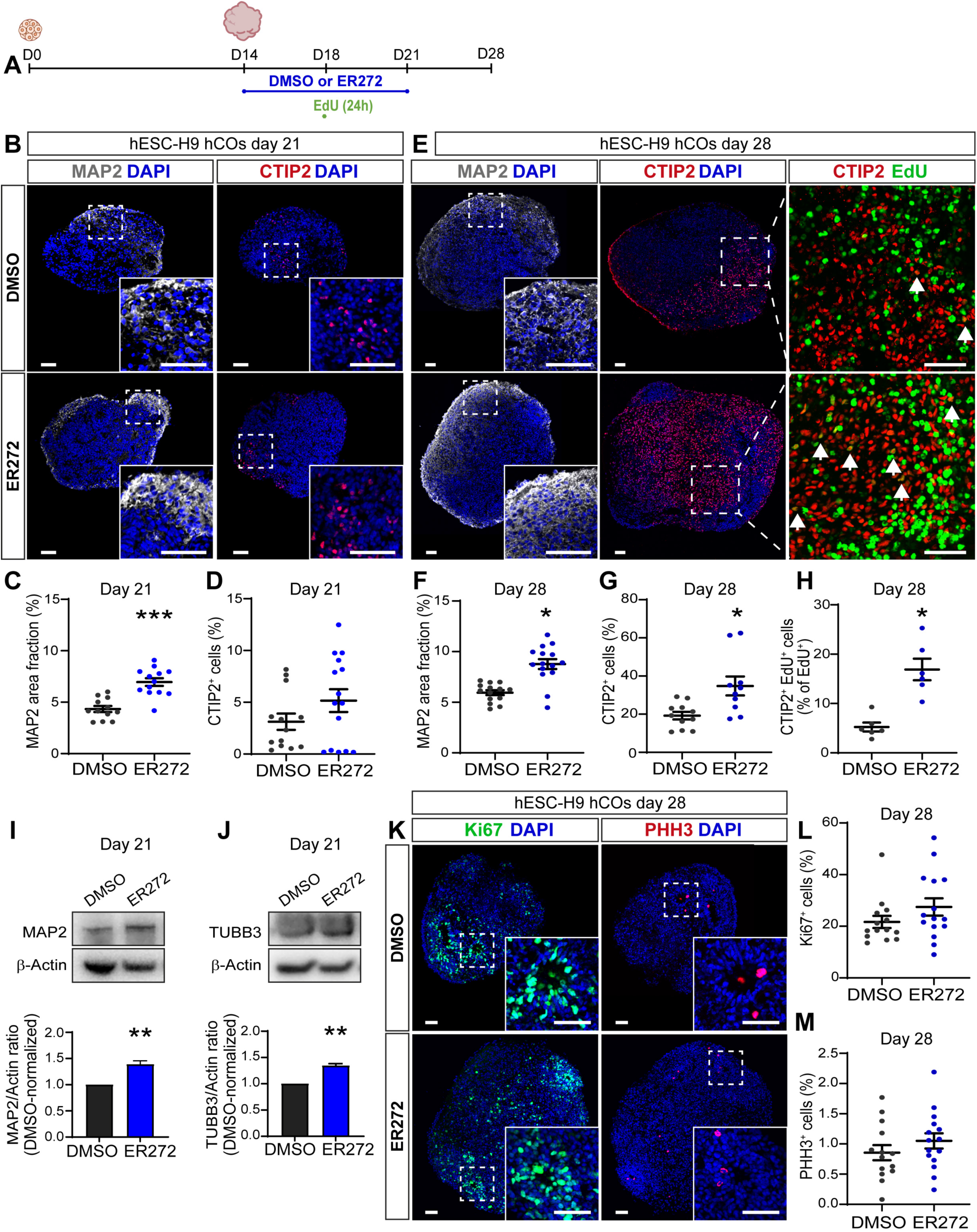
ER272 treatment increases neurogenesis in early-stage human cortical organoids. **(A)** Schematic representation of early-stage H9 hESC-derived hCOs. Organoids were treated with with DMSO (vehicle) or ER272 from day 14 (D14) to day 21 (D21), and a 24h EdU pulse was administered at day 18 (D18). Organoids were collected at D21 or D28 for analysis. **(B)** Images and high magnification confocal views of immunostained organoids for the neuronal marker MAP2 (grey) and cortical layer CTIP2 (red) in day-21 DMSO- or ER272-treated hCOs. Nuclei were counterstained with DAPI (blue). **(C)** Quantification of the percentage of organoid area occupied by MAP2 and **(D)** quantification of the percentage of CTIP2^+^ cells among total DAPI^+^ of cells in DMSO- or ER272-treated H9 hESC-derived hCOs. Values are shown as mean ± SEM from three independent experiments (n=12-15 organoids per condition). **(E)** Images and high magnification confocal views of immunostained organoids for MAP2 (grey), CTIP2 (red), and EdU^+^ cells (green) in day-28 DMSO- or ER272-treated hCOs. **(F)** Quantification of organoid area fraction occupied by MAP2, **(G)** percentage of CTIP2^+^ cells among total DAPI^+^ cells, and **(H)** percentage of CTIP2^+^EdU^+^ cells among total EdU^+^ cells following a 24h EdU pulse and 10-day chase. Values are shown as mean ± SEM from two independent experiments (n=10-15 organoids per condition). For the EdU pulse-chase experiment, data are shown from one experiment (n=6 organoids per condition). **(I, J)** Western blot and quantification of MAP2 and TUBB3, normalized to β-Actin, in day-21 DMSO- or ER272-treated hCOs. Data are mean ± SEM from one experiment (n=3 samples per condition), and expressed relative to DMSO, set 1. **(K)** Immunostaining for the proliferative marker Ki67 (green) and the mitotic marker PHH3 (red) in day-28 hCOs. Nuclei were counterstained with DAPI (blue). **(L, M)** Percentage of Ki67- or PHH3-positive cells among total DAPI^+^ cells in day-28 hCOs. Data are shown as mean ± SEM (n=14-15 per condition, 3 independent experiments). *p < 0.05, **p < 0.01 and ***p < 0.001, Student’s t test. Scale bars, 100 µm.

Next, we tested the effects of ER272 treatment at a later hCO time point in culture which may better represent the mature brain. By contrast, ER272 treatment of late-stage hCOs produced modest or negligible changes in the proportion of SOX2^+^, Ki67^+^, PHH3^+^, EdU^+^, TBR1^+^, CTIP2^+^, and SABT2^+^ cells, as well in MAP2 expression, compared with control hCOs (Fig. S3), suggesting that ER272 has limited pro-neurogenic effects in late-stage hCO cultures.

Overall, these results suggest that ER272 treatment leads to an increase in postmitotic cortical neurons in early-stage hCOs. ER272-induced effects are stage-dependent, with early-stage hCOs displaying widespread changes, whereas late-stage organoids are largely unaffected.

### ER272 promotes early-stage human cortical organoid growth in a PKCα/β1- and Notch-dependent manner

ER272 is a natural reversible C1-domain ligand and activator of the protein kinase C (PKC). In order to understand the mechanism of action of ER272 promoting growth in early hCOs we examined whether Gö6976, a pharmacological inhibitor of PKCα/β1, prevented the effects of ER272 in early-stage hCOs. These experiments showed that whereas ER272 increased total organoid area and number of proliferative and neuronal progenitor cells, this effect was prevented by the PKCα/β1 inhibitor Gö6976 (Fig.3A,C,E-K) in a dose-dependent manner (Fig. S4A,C), implying that ER272 mechanism of action is mediated by PKC α/β1 in hCOs, as suggested before in mouse neurons ^14,15^. Next, we sought to unravel the contribution of the Notch pathway, the master regulator of cell proliferation, on ER272-mediated hCO growth. For this purpose, we used DAPT, a well-known Notch pathway inhibitor which prevents the γ-secretase-mediated cleavage of Notch and the subsequent release of Notch intracellular domain (NICD). Treatment with DAPT reverted the ER272-mediated expansion of the pool of neuronal progenitors and organoid area growth (Fig. 3B,D,E-K and Fig. S4E,G), supporting the role of the Notch pathway.

**Figure 3.**
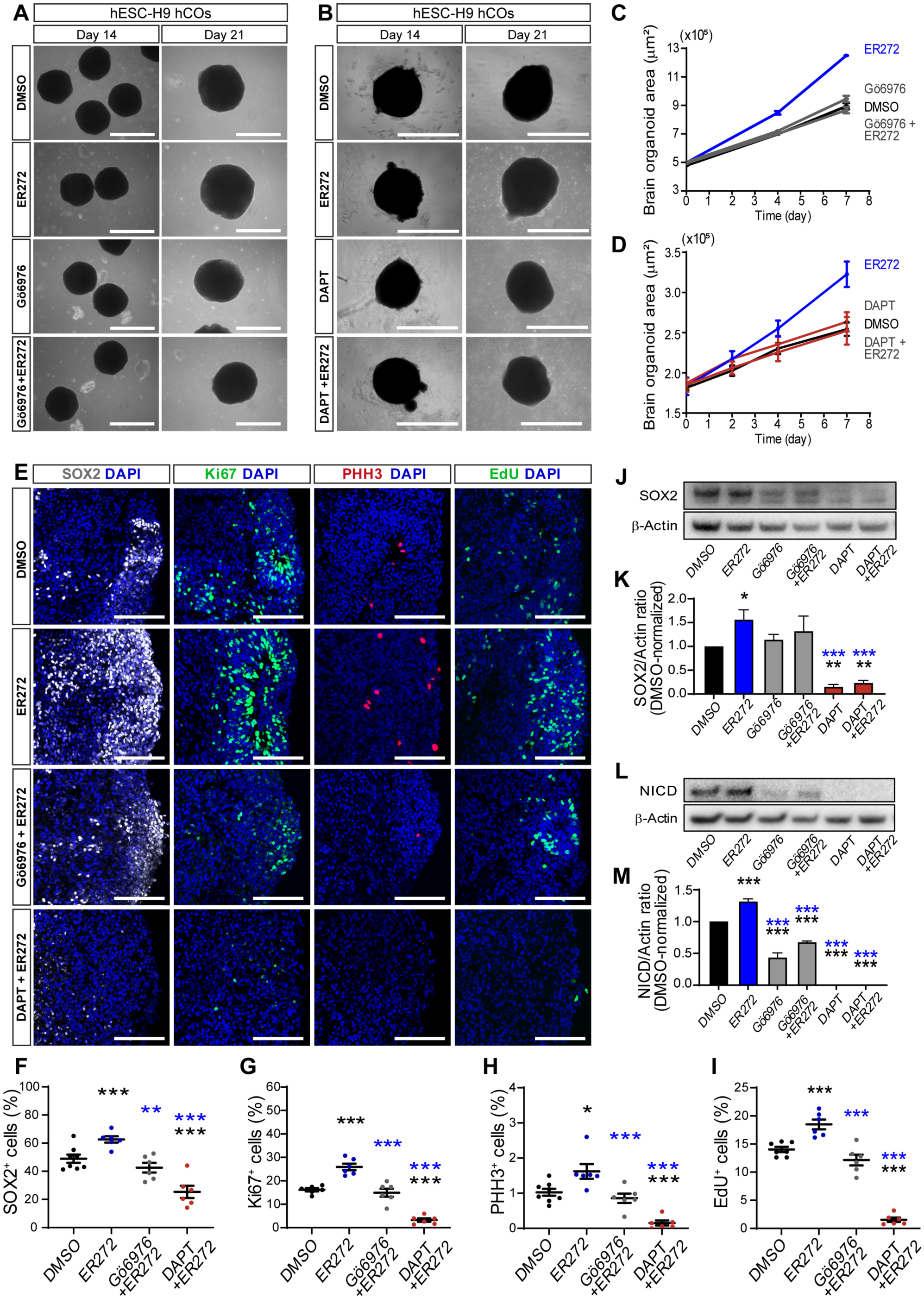
ER272 promotes early-stage human cortical organoid growth in a PKCα/β1- and Notch-dependent manner. **(A-B)** Representative phase-contrast images of live H9 hESC-derived hCOs treated from day 14 to day 21 with DMSO or ER272, in the presence of the classical PKC inhibitor Gö6976 (0.5 µM) or the the γ-secretase inhibitor DAPT (1 µM). Values are shown as mean ± SEM (n=6-10 organoids per condition). **(C-D)** Quantification of organoid area (µm^2^) from day 14 to day 21 following DMSO or ER272, with Gö6976 classical PKC inhibition (C; n = 6-10 organoids per condition) or with DAPT γ-secretase inhibition (D; n = 22-25 organoids per condition). Data are shown as mean ± SEM. ***p < 0.001, two-way ANOVA. **(E)** Confocal views of immunostained organoids for the neural progenitor marker SOX2 (grey), the proliferative marker Ki67 (green), the mitotic marker PHH3 (red) and EdU^+^ (green) cells in day-21 hCOs treated with DMSO or ER272 in the presence or absence of Gö6976 (1 μM) or DAPT (2.5 μM). Nuclei were counterstained with DAPI (blue). **(F-I)** Quantification of the percentage of SOX2-, Ki67-, PHH3- or EdU-positive cells among total DAPI^+^ cells in DMSO or ER272 treated hESC-H9 hCOs in the presence of the classical PKC inhibitor Gö6976 or γ-secretase inhibitor DAPT (n=6 organoids per condition). **(J-K)** Western blot and quantification of SOX2 protein levels normalized to β-actin (n = 3 samples per condition). **(L-M)** Western blot and quantification of NICD protein levels normalized to β-actin (n = 3 samples per condition). Protein levels are expressed relative to DMSO, set to 1. Data are shown as mean ± SEM. Black asterisks indicate comparisons with DMSO and blue asterisks comparisons with ER272. *p < 0.05, **p < 0.01, and ***p < 0.001, one-way ANOVA. Scale bars, 1000 µm for phase-contrast images; and 100 µm for immunofluorescence images.

Given that the PKCα/β1 and Notch pathways mediate ER272-driven organoid growth, we sought to determine if they also regulate basal hCO growth in the absence of drug treatment. To answer this question, we performed a dose curve experiment using either Gö6976 or DAPT in early hCOs in culture. Strikingly, we found that Gö6976 treatment led to basal hCO growth inhibition in a dose-dependent manner (Fig. S4B,D), similarly to the use of DAPT (Fig. S4F,H). Further, direct assessment of the NICD levels, the effector of Notch signaling, following ER272-treatment showed an increase compared to control hCOs, indicative of higher Notch activity (Fig. 3L-M). Conversely, the treatment of hCOs in basal conditions with the PKCα/β1 inhibitor Gö6976 showed a drastic decrease in the levels of NICD, suggesting lower Notch activity. Interestingly, and in agreement with our previous findings, the combination of ER272 and Gö6976 reverted the NICD increase produced by ER272 treatment. As a control we showed that DAPT treatment led to complete NICD depletion. Finally, a combination of DAPT and ER272 also reverted the NICD increase following ER272 treatment (Fig. 3L-M).

All these results indicate that basal and ER272-induced hCO growth requires PKCα/β1 and Notch signaling and suggest that PKC activity contributes to the regulation of NICD levels.

### ER272 treatment upregulates extracellular matrix, cell adhesion and growth factor-related genes in early-stage human cortical organoids

To evaluate the extent of the overall transcriptomic changes triggered by ER272 treatment in hCOs we performed bulk RNA sequencing of control (DMSO) and ER272-treated hCOs at day 21. Principal component analysis (PCA) showed a separation between control and ER272-treated samples along PC1 (Fig. 4A). We found 410 genes upregulated and 152 genes downregulated in ER272-treated hCOs when compared to control hCOs (DMSO) (Fig. 4B, Supplementary Table 1). Among the top 15 upregulated genes we identified genes related to growth and proliferation such as ceruloplasmin (*CP*); extracellular matrix organization, metalloprotease activity and cell adhesion including adhesion G protein-coupled receptor L4 (*ADGRL4*), ADAM metallopeptidase domain 12 (*ADAM12*), and intercellular adhesion molecule 1 (*ICAM1*); as well as signal transduction related genes, such as solute carrier family 5 member 7 (*SLC5A7*) (Fig. 4C). Consistent with these changes, enrichment pathway analysis of the upregulated DEGs identified extracellular matrix, signal transduction, and receptor tyrosine kinase-related pathways among the top 10 most enriched pathways using Reactome database (Fig. 4D, Supplementary Table 2). Among those, we observed increased expression of tyrosine kinases and growth factor/proliferative genes next to extracellular matrix genes, cell adhesion molecules and integrins (Fig. 4E-F and Fig. S5). Interestingly, genes encoding PKC isoforms (*PRKCA, PRKCD*), epidermal growth factor (*EGF*), ligand of the EGFR, the receptor tyrosine kinase *ERBB3*, the transcription factor *STAT3* which promotes cell proliferation, as well as cyclins and cyclin-dependent kinases (*CCNA1*, *CDKN1A* and *CDKN1C)* were upregulated in ER272-treated hCOs (Fig. 4F), suggesting an increase in the expression of the PKC enzymes, the EGFR ligand and EGFR, and in genes implicated in regulation of the cell cycle and cell proliferation. Moreover, we found *ADAM12*, *ADAMTS3*, *ADAMTS4*, members of the metalloprotease ADAM family, CDH19, ICAM1, ITGAV, ITGB5, cell adhesion and integrin genes, as well as the downstream Notch targets *FABP7*, *HEYL,* upregulated in hCOs following ER272 treatment (Fig. 4F and Fig. S5), suggesting changes in the expression of enzymes that process extracellular matrix components and that may modulate Notch processing, next to changes in the expression of cell adhesion molecules and downstream targets of Notch signaling.

**Figure 4.**
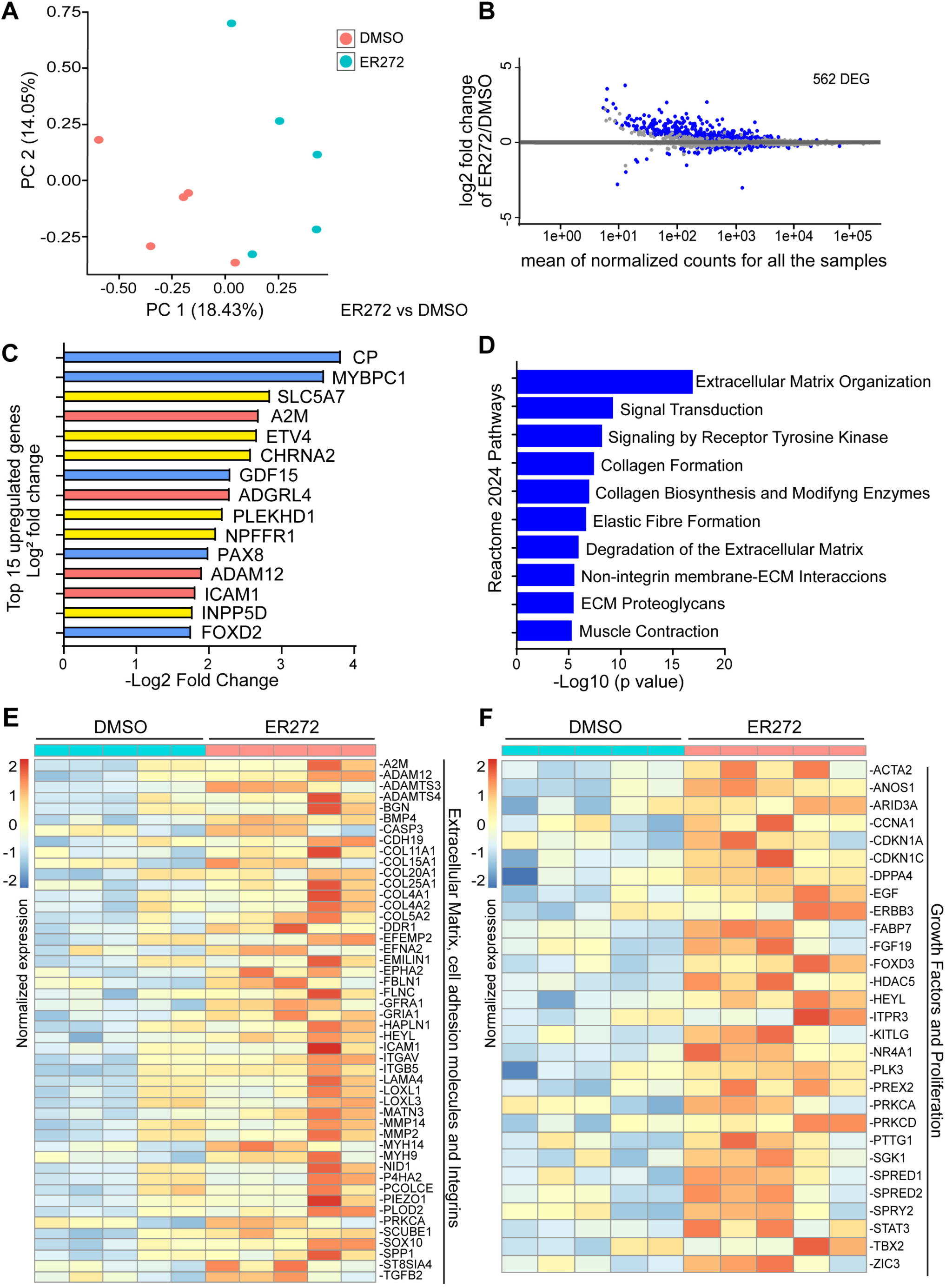
ER272 treatment upregulates extracellular matrix, cell adhesion and growth factor-related genes in early-stage human cortical organoids. **(A)** Principal component analysis (PCA) of bulk-RNA-seq data from DMSO- and ER272-treated hCOs at day 21 (two independent experiments, n = 5 organoids per condition), showing the different dynamics of the different samples. The x-axis represents the principal component 1 (PC1) that separates with 18.43% of the variance the DMSO and ER272 treated hCO samples. **(B)** MA plot showing the 562 differentially expressed genes (DEGs) in ER272-treated hCOs compared with DMSO controls (410 upregulated and 152 downregulated). Values represent the Log2 fold change on the y -axis (M) and the mean of normalized counts for all samples on the x -axis (A). Genes with adjusted *p* < 0.05 are shown in blue; non-significant genes are shown in grey. **(C)** Top15 upregulated DEGs based on the Log2 fold change from ER272 treated hESC-H9 hCOs compared to DMSO treatment. In blue: growth and proliferation related genes; in red: metalloproteases and integrins genes; in yellow: signal transduction related genes. **(D)** Pathway enrichment analysis of upregulated DEGs comparing ER272- and DMSO-treated hCOs. The top 10 Reactome 2024 pathways are shown. The x - axis represents the -log10 p-value of pathway enrichment. **(E-F)** Heatmaps of the upregulated genes associated with Extracellular matrix, cell adhesion and integrin (E) and Growth factors and proliferation (F) pathways in ER272-treated hCOs compared with DMSO-treated hCOs. VST-transformed read counts are shown, with each row representing one gene and each column one independent hCO sample.

Collectively, all these data suggest that ER272 treatment in early-stage hCOs leads to transcriptomic changes related to extracellular matrix, tyrosine kinase and growth factor genes, consistent with a role of the PKCα/β1 and Notch pathways.

### OGD-induced ischemic-like injury in late-stage human cortical organoids is not rescued by ER272 treatment alone

To model ischemic-like injury *in vitro* and evaluate the regenerative potential of ER272 following injury, late-stage hCOs derived from hiPSC were exposed to oxygen-glucose deprivation (OGD) for 6 h and subsequently treated for 7 days with either DMSO or ER272 (Fig 5A). OGD increased the proportion of cleaved caspase-3⁺ (C-CASP3) cell compared with control hCOs, indicating increased apoptotic cell death in ischemic-like conditions *in vitro* (Fig. 5B-C). Despite this increase in apoptosis, OGD did not affect total organoid area or cell density, implying discrete effects of OGD-induced ischemia-like conditions in hCOs (Fig. S6A-C). OGD increased both GFAP⁺ area and the proportion of Ki67⁺ cell compared with controls, consistent with an astroglial response and increased cell proliferation after ischemic-like injury (Fig. 5B,D and Fig. S6A,D), as shown before *in vivo* ^44–46^. We next analyzed the changes produced by OGD in the neuronal population in late-stage hCOs. We found reduced proportion of the pan-neuronal markers HuC/D, as wel as reduced expression of MAP2 and TUBB3, next to reduced proportion of CTIP2 cortical cells, suggesting the loss of neurons following ischemic-like conditions *in vitro* (Fig. 5B,E-F and Fig. S6A,E-F). Finally, we evaluated the effects of ER272 treatment following OGD in hCOs. Notably, ER272 treatment alone did not reduce the proportion of C-CASP3⁺ cells or rescued OGD-induced changes in GFAP⁺ area or the proportion of Ki67⁺ cell, which both remained elevated compared with controls (Fig. 5B-D and Fig. S6A,D). Moreover, ER272 treatment was not sufficient to rescue the loss of neuronal markers under OGD conditions in late-stage hCOs (Fig. 5B,E-F and Fig. S6A,E-F).

**Figure 5.**
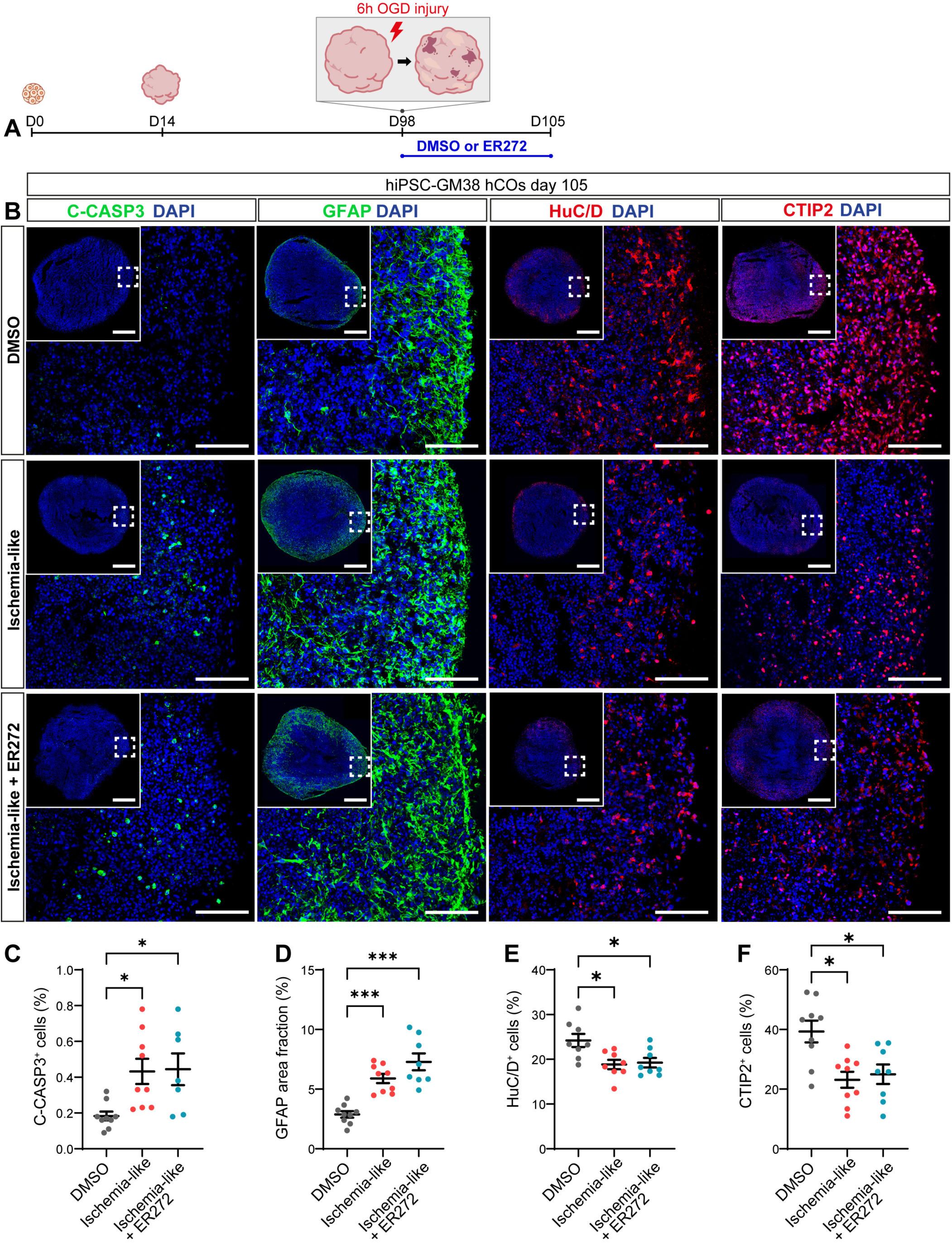
OGD-induced ischemic-like injury in late-stage human cortical organoids is not rescued by ER272 treatment alone. **(A)** Experimental design for GM38 hiPSC-derived late-stage hCOs. At day 98 (D98), organoids were exposed to 6 h of oxygen-glucose deprivation (OGD) to induce an ischemia-like injury or maintained under normoxic control conditions. Following OGD, organoids were treated by 1 week of DMSO or ER272 treatment. **(B)** Representative tile scan and high magnification confocal views of immunostained organoids for cleaved caspase-3 (CASP3; green), GFAP (green), HuC/D (red), and CTIP2 (red), in control, ischemia-like, and ischemia-like + ER272 hCOs in late-stage hCOs. Nuclei were counterstained with DAPI (blue). Insets show low-magnification images indicating the imaged region. **(C-F)** Quantification of the percentage of CASP3-positive cells, GFAP-positive area fraction, HuC/D-positive cells, and CTIP2-positive cells, respectively. Data are shown as mean ± SEM (n = 8-10 organoids per condition). *p < 0.05 and ***p < 0.001, one-way ANOVA. Scale bar, 100 μm, insets 500 μm.

Together, these results show that OGD recapitulates features of ischemia-like injury in late-stage hCOs, including apoptosis, a proliferative response, astroglial reactivity, and loss of human neurons. In addition, consistent with the lack of effect observed in mature organoids, ER272 did not significantly rescue the main neuronal alterations induced by OGD under these conditions.

### Organoid-to-organoid transplantation combined with ER272 treatment enhances graft integration and proliferation after ischemia-like injury

Given that ER272 treatment alone did not rescue the main alterations induced by ischemia-like injury in mature hCOs, we next tested whether its effects on early-stage hCOs could be exploited in an organoid-to-organoid transplantation paradigm following OGD. Early-stage GFP⁺ hCOs were transplanted onto mature host ischemic-like hCOs immediately after OGD and treated for 7 days with either DMSO or ER272 (Fig 6A-B). ER272 treatment expanded the GFP⁺ graft area (Fig. 6C,E), consistent with enhanced early-stage hCO growth. Accordingly, we found a higher proportion of Ki67⁺GFP^+^ cells within the graft following ER272 treatment under OGD ischemia-like conditions (Fig. 6C,F), implying a proliferative effect of the compound on grafted early-stage hCOs. Given that ER272 treatment promoted the expression of extracellular matrix genes, cell adhesion molecules and integrins in early-stage hCOs, we evaluated the extent of integration of the graft into the host ischemic-like injured hCO tissue following ER272 treatment. The evolution of the transplanted early GFP⁺ hCO was monitored by live brightfield and GFP fluorescence imaging (Fig. 6D). At 7 days post-transplantation (7 DPT), GFP⁺ grafts remained predominantly positioned at the host surface in control and OGD ischemia-like hCOs (Fig. 6D). Remarkably, ER272 treatment in ischemia-like hCOs significantly improved the integration of the graft into the host organoid (∼90% GFP^+^ graft integrated) when compared to control or OGD conditions which remained similar (∼40% GFP^+^ graft integrated) (Fig. 6D,G).

**Figure 6.**
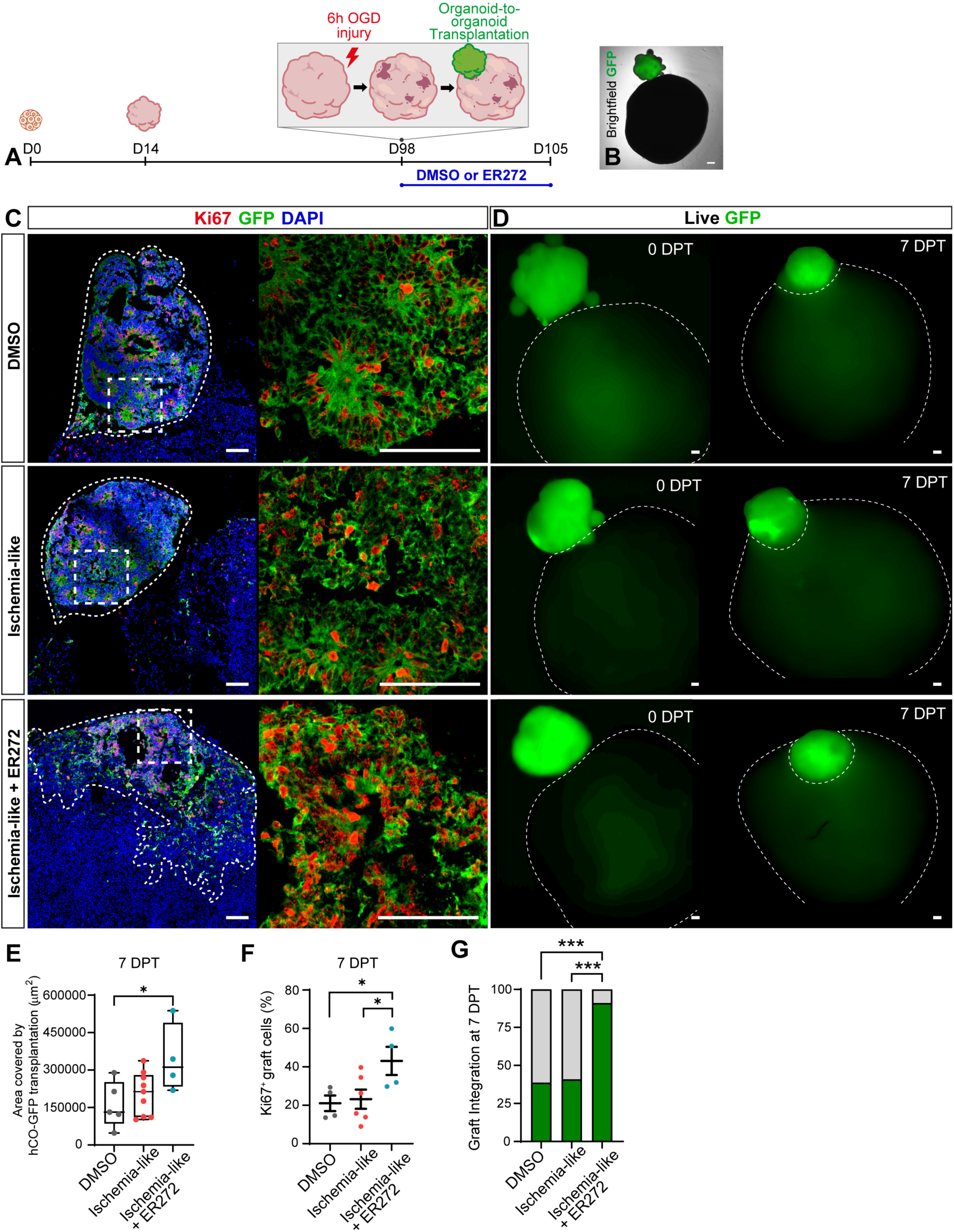
Organoid-to-organoid transplantation combined with ER272 treatment enhances graft integration and proliferation after ischemia-like injury. **(A)** Schematic representation of the experimental design for GM38 hiPSC-derived human cortical organoids (hCO). At day 98 (D98), host late-stage hCOs were subjected to 6 h of oxygen-glucose deprivation (OGD) to induce an ischemia-like injury, followed by transplantation of GFP-labeled donor early-stage hCOs. After transplantation, hCOs were treated with DMSO (vehicle) or ER272 for 7 days and analyzed at 7 days post-transplantation (7 DPT). **(B)** Representative brightfield and GFP fluorescence image showing a GFP-labeled donor early-stage hCO attached to a host late-stage hCO following organoid-to-organoid transplantation. **(C)** Confocal views of immunostained organoids for Ki67 (red) and GFP (green) in DMSO, ischemia-like, and ischemia-like + ER272 hCOs at 7 DPT. Insets show higher magnification of the boxed regions. **(D)** Representative live GFP images at 0 and 7 DPT in DMSO, ischemia-like, and ischemia-like + ER272 hCOs at 0 and 7 DPT. Nuclei were counterstained with DAPI (blue). **(E)** Quantification of the GFP-positive graft area (μm^2^) at 7 DPT. **(F)** Percentage of proliferating GFP^+^ graft cells, expressed as Ki67^+^ GFP^+^ cells among total DAPI^+^GFP^+^ cells at 7 DPT. **(G)** Quantification of graft integration at 7 DPT, expressed as the percentage of the GFP-positive graft area located within the host organoid. Data are shown as mean ± SEM (n = 4-6 organoids per condition, 2 independent experiments). ***p < 0.001, one-way ANOVA. Scale bars, 100 μm.

These results show that dual transplantation and ER272 treatment leads to graft proliferation and enhanced integration into OGD-injured late-stage hCOs *in vitro*, consistent with the previously observed ER272-mediated effects in early-stage hCOs.

### Combined cell transplantation and ER272 pro-neurogenic treatment rescue neuronal loss in ischemic-like late-stage human cortical organoids

Since ER272 promotes proliferation and subsequent neuronal differentiation in early-stage hCOs, we hypothesized that ER272 treatment of early hCO grafts could enhance their neurogenic response following transplantation into OGD-injured late-stage hCOs. To test this, transplanted hCOs were treated with DMSO or ER272 for 7 days after transplantation and maintained for an additional 7 days without treatment, allowing time for proliferating cells to undergo neuronal differentiation. At 14 days post-transplantation (14 DPT), GFP⁺ graft area showed a tendency to increase in the ER272-treated group (Fig. 7A-B), consistent with the results observed at 7 DPT. Similarly, ER272 treatment markedly enhanced graft-host integration at 14 DPT (Fig. 7A, C). Notably, ER272 treatment also increased neuronal marker expression within the GFP⁺ graft, including the pan-neuronal marker HuC/D, the cortical marker CTIP2, and the neuronal marker MAP2 (Fig. 7A, D–F). Importantly, this increase in neuronal markers within the graft was accompanied by a recovery of the loss of neurons observed under ischemic-like conditions at the whole organoid-to-organoid level to control hCO levels (Fig. 7A, G-I). In addition, we found that ER272-treated organoids showed increased number of GFP^+^ cells and neuronal fibers, further supporting enhanced graft-host integration and reestablishment of lost neuronal tracts under ischemic-like conditions (Fig. 7J).

**Figure 7.**
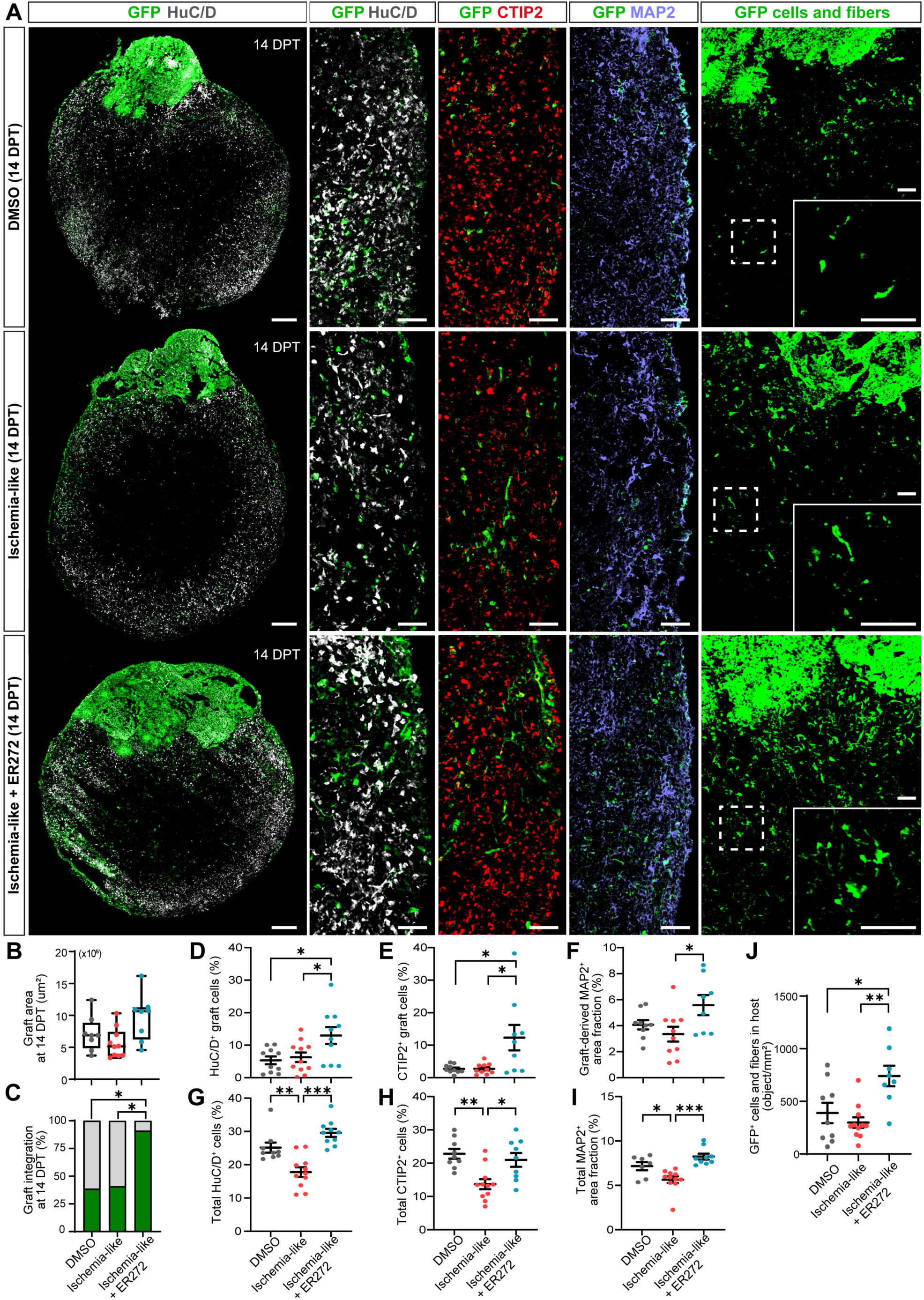
Combined cell transplantation and ER272 pro-neurogenic treatment rescue neuronal loss in ischemic-like late-stage human cortical organoids. **(A)** Representative tile scan and high magnification confocal views of immunostained organoid-to-organoid transplantation model under three experimental conditions: vehicle-treated control (DMSO), ischemia-like injury, and ischemia-like injury treated with ER272 at 14 days post-transplantation (14 DPT). Donor-derived graft cells are identified by GFP (green), while neurons are labeled with HuC/D (gray), deep-layer cortical neurons with CTIP2 (red), and mature neurons with MAP2 (purple). Insets highlight representative GFP⁺ cells and fibers. **(B)** Quantification of the total GFP^+^ graft area (µm²) at 14 DPT. **(C)** Percentage of graft integration within the host organoid, expressed as the percentage of the GFP^+^ graft area located within the host organoid. **(D-F)** Percentage of HuC/D⁺ cells, CTIP2⁺ cells, and MAP2⁺ area fraction (%) within the graft GFP^+^ area (HuC/D⁺GFP^+^ among DAPI⁺GFP^+^, CTIP2⁺GFP^+^ among DAPI⁺GFP^+^, and MAP2⁺GFP^+^ area fraction respectively). **(G-I)** Percentage of total HuC/D⁺ cells (HuC/D⁺ among DAPI⁺); CTIP2⁺ cells (CTIP2⁺/DAPI⁺); and MAP2⁺ area fraction (%) in the whole organoid-to-organoid transplantation model. **(J)** Number of GFP⁺ graft-derived cells and GFP^+^ fibers within the host organoid. Data are presented as mean ± SEM (n = 8-11 organoids per condition, 2 independent experiments). Each dot represents one organoid. Statistical significance was determined by one-way ANOVA with Tukey’s multiple comparisons test. \**P* < 0.05, \*\**P* < 0.01, \*\**P* < 0.001. Scale bars: low-magnification images, 200 μm; high-magnification images, 50 μm; insets, 50 μm.

Together, these results show that dual organoid transplantation and ER272 treatment promote graft integration and recovery of neuronal loss in ischemic-like late-stage hCOs *in vitro*.

## DISCUSSION

Cell transplantation therapies and pro-neurogenic compound treatments constitute alternative strategies for brain repair, which are undergoing clinical trials for the treatment of cognitive impairment, Parkinson’s disease, Alzheimer’s disease and ischemic stroke patients. Although both approaches have shown regenerative potential in preclinical models, their translation into effective and durable treatments for the human brain remains challenging ^47^.

Interestingly, recent preclinical studies have introduced the use of human brain organoids for *in vivo* transplantation as an alternative to conventional transplantation of neuronal dissociated cells ^48–50^. These studies have highlighted the advantages of 3D human brain organoids as a donor, revealing a higher graft cell diversity that includes neurons, and glial cells, improved long-term cell survival following transplantation, and displaying a higher neuronal recovery rate within the damaged mouse brain ^51–55^.

Here, we show that ER272 pro-neurogenic treatment induces brain organoid growth through PKC and Notch-dependent pathways, ultimately increasing the production of cortical neurons in early-stage hCOs. We provide a molecular analysis of early-stage hCOs treated with ER272 and found upregulation of growth factors, integrins, cell adhesion molecules and extracellular matrix related genes. We show that ER272 treatment combined with early-stage hCO transplantation into ischemic-like late-stage hCOs results in increased integration and graft growth into the host, that led to the replacement of the lost neurons and neuronal fibers.

This work brings to the field several novel aspects. First, we show for the first time that ER272 treatment of human cortical organoids leads to an expansion of the neural progenitor pool promoting organoid growth, that results in increased cortical neuronal number. Interestingly, this effect was hCO stage-dependent: it was robustly displayed by early-stage hCOs, which may reflect an embryonic-like neurogenic environment, but was largely absent or significantly reduced in late-stage hCOs *in vitro*, which better mimic aspects of a more mature brain state. This discrepancy suggests that early hCOs provide a receptive, embryonic-like cellular context for pro-neurogenic signals like ER272, whereas late mature hCOs present a more restrictive state that limits those responses. In agreement with this, we detected a higher number of progenitor cells and cells in proliferation in hCOs in basal conditions at day 21 (early stage; 20% Ki67^+^ cells) than at day 105 *in vitro* (late stage; 5% of Ki67^+^ cells). As a result, ER272 treatment promoted cortical neurogenesis in early hCOs (2-fold and 3.5-fold increase in total and newly born CTIP2^+^ cells, respectively), but its effect was negligible in late-stage hCOs. This difference in drug-response pattern reflects that early-stage hCOs may represent an immature stage similar to the early embryonic brain, whereas late-stage hCOs may parallel more mature human neural tissue devoid of cells in proliferation. Our work also highlights intriguing differences between human and murine cells. While previous reports showed that ER272 treatment stimulates endogenous stem cell proliferation and neurogenesis in the SAMP8 aged mouse brain ^15^, our late-stage hCOs did not exhibit this same robust response. This discrepancy likely reflects fundamental species-specific differences in how mature human and mouse tissues respond to pro-neurogenic signals. Alternatively, it is possible that cortical organoids lack the specific neurogenic niches of the adult brain, such as the subventricular zone (SVZ) or the dentate gyrus (DG) of the hippocampus, which are known to support the neurogenic responses elicited by ER272 in the adult murine brain. Hence, future studies leveraging region-specific brain organoids containing SVZ or SGZ niches will be valuable to test whether human late-stage organoids *in vitro* can support ER272-mediated responses.

Second, our data show that, alongside the well-known Notch signaling pathway, PKCs also play a role in the proliferation of early-stage hCOs under basal conditions in the absence of ER272 treatment. Further, we observed a reduction in NICD levels following PKCα/β1 inhibition, suggesting a role for PKC signaling in Notch pathway regulation in early-stage hCOs. This is consistent with previous studies showing that atypical PKC isoforms, such as PKCζ and PKCι, regulate Notch signaling, supporting a functional crosstalk between PKC and Notch pathways ^56,57^. However, to our knowledge, a functional link involving conventional PKC isoforms (α/β1) and Notch in regulating neuronal proliferation in the human embryonic cortex has not been previously established. Future studies will be essential to further elucidate the underlying molecular mechanisms driving this crosstalk.

Third, we observed that ER272 alters the expression profile of growth factors, integrins, cell adhesion molecules and extracellular matrix genes in early-stage hCOs. Although prior work demonstrated that ER272 activates classic PKCs and induces the release of growth factor ligands ^14,15^, its downstream transcriptomic consequences in neural cells remained unexplored. Our results confirm that ER272 pro-neurogenic treatment triggers changes in signaling pathways in brain cells related to cell proliferation through upregulation of PKC, PKA, EGF, ERBB3, but also of growth factors and secreted proteins like TGFB2 and BMP4, cyclins and cyclin-dependent genes such as CCNA1, CDKN1A and CDKN1C. Moreover, our results show that ER272 treatment upregulates the expression of integrins such as ITGAV, ITGB5, cell adhesion molecules such as ICAM1, CDH19, next to the expression of matrix metalloproteinases and proteases such as MMP2, MMP14, ADAM12, ADAMTS3, ADAMTS4 and various collagen chain genes, highlighting an important role in remodeling and sensing the extracellular matrix that may also be relevant to the cellular processes underlying graft integration.

Fourth, we introduce a novel human *in vitro* organoid-to-organoid transplantation model to analyze the potential of dual strategies combining pro-neurogenic treatment with stem cell-derived transplantation for the recovery of the post-ischemic stroke brain. This platform could serve as a screening tool for combinatorial drug treatments targeting cell loss and/or for performing compound dose-response analyses within a human relevant context. Lastly, our study reveals that ER272 combined with early-stage human organoid transplantation has the potential to display the full effect of a pro-neurogenic treatment resulting in higher neuronal counts, and further, we show for the first time that ER272 improves transplant integration into the host and increases transplant outgrowth, supporting further investigation of combined pharmacological and transplantation-based strategies for brain repair.

Previous *in vivo* xenotransplantation work showed that hiPSC-derived cortical cells establish local and long-range axonal projection patterns with the host comparable to those of native cortical neurons, emphasizing their ability to selectively reconstruct cortical tracts within the lesioned adult mouse brain ^1,2,26,28^. A limitation of our current model is the lack of multi-region organoid structures to investigate connectivity patterns following specific grafts and treatments. Future improvements of our *in vitro* system could integrate assembloids to model multi-region human brain architecture, enabling the analysis of axonal projection specificity following transplantation and pro-neurogenic interventions.

We and others have shown that *in vivo* cell transplantation requires an identity match between the damaged area and the donor cells in order to display its full regenerative potential within the adult mouse brain ^1,26,58^. These studies demonstrated that transplantation of midbrain dopamine neurons or glutamatergic neurons within the substantia nigra (SN) of Parkinson’s disease (PD) mouse models resulted in different outcomes, the former efficiently innervating the dorsal striatum, whereas the latter failed to project axons to the SN ^58^. Within the lesioned cortex, the transplantation of non-matching cortical areal identity cells led to failure of graft integration and innervation of cortical and subcortical-specific targets, in contrast with the reestablishment of cortical tracts following matched graft areal identity ^1,5,26^. Future studies using our human organoid-to-organoid transplantation platform could further assess the limitation of non-matched grafted cells, such as thalamic, midbrain or cerebellar organoids, to integrate and recover cortical neuronal loss following ischemic-like conditions in hCOs.

Reconstruction of regional-specific axonal tracts requires the presence of receptors and ligands acting in coordination to guide the growth of axons and to direct synapse formation. Several studies have investigated the secreted signals and factors that may still be present or could be re-expressed following a lesion of the adult brain and that promote axogenesis following transplantation ^59–63^. Interestingly, neurexin, neuregulin, neural cell adhesion molecule and SLIT have been shown to be involved in graft-host crosstalk ^59,60^. In this respect, it would be interesting in the future to investigate whether these signals are present in our organoid-to-organoid ischemic transplantation paradigm, and whether a combination with pro-neurogenic drugs may further potentiate these effects.

Lastly, stem cell transplantation into the damaged mouse brain has been shown to reduce endogenous astrocyte and microglia activation, reducing neuroinflammation associated with improved post-stroke outcomes ^59,64–66^. Future work should develop an immunocompetent organoid-to-organoid transplantation model for ischemia that includes the presence of hiPSC-derived microglia cells. This model could be instrumental to understand the contribution of human astrocytes and human microglia to the downstream effects provoked by stroke and their contribution to post-stroke recovery pathways.

*In vitro* 3D human brain organoid disease models offer unique opportunities for the study of human-specific pathological features and to evaluate effective treatments. Our work establishes a novel *in vitro* human organoid-to-organoid ischemic-like transplantation model to study novel mechanisms that can help repair the adult human brain. Our findings support that a combined stem cell transplantation therapy and pro-neurogenic treatment can improve the outcome of transplantation alone by facilitating integration into the host tissue and promoting graft neural growth. Altogether, these findings highlight that the use of combined therapeutic approaches may be potentially useful for the field of brain regeneration.

## RESOURCE AVAILABILITY

Further information and requests for resources and reagents should be directed to, and will be fulfilled by, the lead contact, Ira Espuny-Camacho.

RNA-seq analysis codes can be found in the following repository: https://github.com/Espuny-Camacho/GOMEZ-OLIVA_2026/

## ACKNOWLEDGEMENTS

We thank GIGA Imaging, GIGA Genomics and Bioinformatics platforms for their contribution, help and support to this work.

This work was supported by the Credit Classique 2020 from the Fonds Léon Fredericq and the Faculty of Medicine from the University of Liège (to I.E.-C.); the Credit Sèctoriel, Fonds Speciaux 2024 from the University of Liège (to I.E.-C.); the European EMBO-Short Term Fellowship (to R.G.-O.); the Erasmus+ European Mobility KA103 (to R.G.-O.); and the Plan Propio UCA 2022-2023 program from the University of Cádiz, Spain (to R.G.-O.).

## AUTHOR CONTRIBUTIONS

I.E-C designed the initial study. C.C. and R.H-G synthesized the diterpenes for this study. R.G-O, designed the experimental plan and performed the experiments with the help of the rest of co-authors. S.H. performed the bioinformatics analysis. R.G-O analyzed all the results. I.E-C and R.G-O wrote the manuscript with the help of the rest of coauthors.

## SUPPLEMENTAL INFORMATION

Document S1. Supplemental methods and Figures S1-S6

Large table S1. Differentially expressed genes (DEGs).

Large table S2. Reactome Pathways 2024 enrichment analysis.

## MATERIALS AND METHODS

### Human cortical organoid (hCO) differentiation

Cortical brain organoids were developed following a modified version of the Sasai protocol ^33^. hPSCs were dissociated and seeded into 96 well U bottom plates at 9,000 cells per well. Organoids were cultured in DMEM-F12 (ThermoFisher 31331-093) with 20% KO serum (Fisher 10828028), 1/100 non-essential amino acid (NEAA; ThermoFisher 11140-035), 1/100 penicillin/streptavidin (Pen/strep; ThermoFisher 15140122) and 100µM β-mercaptoethanol (Sigma 444203) for 14 days. 5µM SB (SB431542; Sigma S4317) and 3µM IWR1 (Sigma ref: 681669) were added from day 0-14. Rock inhibitor (Y-27632 2HC; Bio-Connect S1049) was added on day 0 at 20µM and at 10µM from day 2 to day 6. From day 14 to day 28, medium was composed of DMEM-F12, 1/100 sodium pyruvate (Na Pyr; ThermoFisher 11360-039), 1/100 NEAA, 1/100 N2 (ThermoFisher 17502048), 1/100 Pen/strep, 1/150 BSA (BSA fraction V; ThermoFisher 9048-46-8), 1/50 B27 without vitamin A (ThermoFisher 11500446) and 100µM β-mercaptoethanol. From day 28 to day 35, medium was composed of half DMEM-F12 and half Neurobasal (ThermoFisher 21103-049) 1/200 Na Pyr, 1/200 NEAA, 1/200 N2, 1/200 Glutamax (ThermoFisher 35050-038), 1/300 BSA, 1/100 Pen/strep, 1/50 B27 without vitamin A and 50µM β-mercaptoethanol. From day 35 we used B27 with vitamin A (ThermoFisher 11530536). At day 70, 1% of Matrigel (Corning 354230) was added to the culture.

### Organoid-to-organoid transplantation model

An organoid-to-organoid transplantation model was established by gently positioning 2-week-old GFP-labelled early cortical organoids (hCOs) in contact with the surface of 98-day-old late-stage hCOs for 7 days or 14 days *in vitro*.

### RNA isolation and sequencing, data analysis and differential expression analysis

RNA was extracted using the RNeasy mini kit (Qiagen 74104), followed by a DNase treatment (Ambion AM2222). High-quality RNA samples, (RIN) values of 8-10, were used for sequencing. hESC-H9 hCOs treated with DMSO (n=5) or treated with ER272 (n=5) from 2 independent differentiations were analyzed. PolyA RNA-seq libraries (Illumina) were sequenced using the Illumina Novaseq 6000 sequencer, generating 2 x 150-bp paired-end reads. Downstream analysis was performed in R using DESeq2 method for normalization, clustering and differential expression analysis.

### Statistical analysis

All statistical analysis was performed using GraphPad (Prism). For comparisons between two groups, an unpaired, two-tailed Student’s *t*-test was used for normally distributed data, whereas the corresponding non-parametric Mann– Whitney *U* test was used when the data did not follow a normal distribution. For comparisons involving more than two groups, one-way ANOVA followed by Tukey’s multiple-comparisons test was used for normally distributed data, whereas the Kruskal-Wallis test followed by Dunn’s multiple-comparisons test was used for non-normally distributed data. For experiments involving two independent variables, two-way ANOVA followed by Šídák’s multiple-comparisons test was used. The specific statistical test, and sample size (*n*) for each analysis are indicated in the corresponding figure legends. Statistical significance was defined as *p* < 0.05; p < 0.01; *p* < 0.001.

