## Supplemental materials for "Organoid-to-organoid transplantation and pro-neurogenic treatment promote graft integration and neuronal repair in a human ischemic model"

### Supplemental Figures

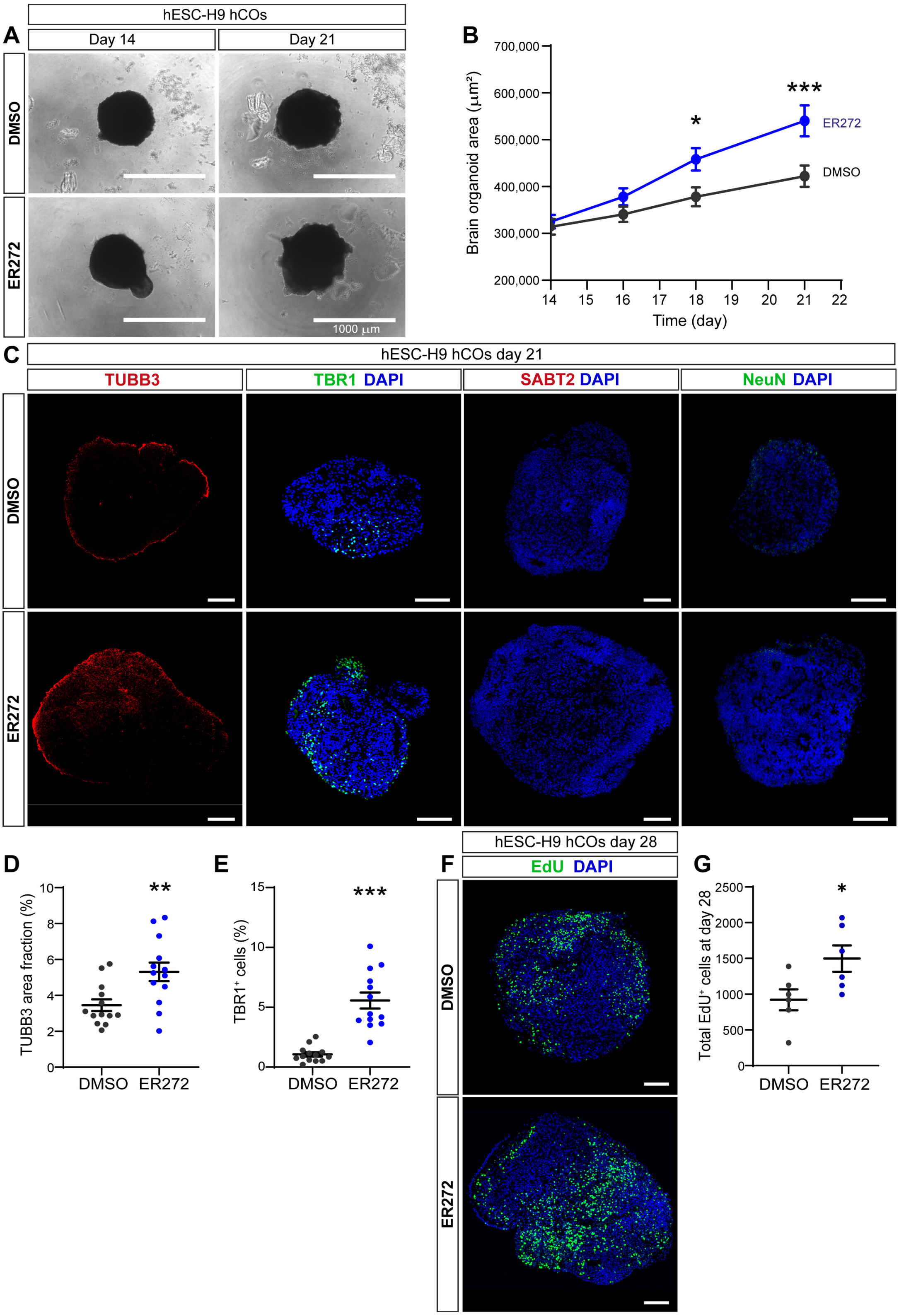


**Supplementary Figure 1. ER272 enhances growth, neuronal differentiation, and proliferation in human cortical organoids (hCOs).** **(A)** Bright-field images of H9 human embryonic stem cell-derived cortical organoids (hESC-H9 hCOs) cultured in the presence of vehicle (DMSO) or ER272 from day 14 to day 21. Scale bars, 1000 μm. **(B)** Quantification of hCO area during the treatment period (days 14-21). **(C)** Representative immunofluorescence images of day 21 hCOs stained for the deep-layer cortical neuron marker TBR1 (green), the neuronal marker βIII-tubulin (red), the upper-layer cortical neuron marker SATB2 (red), and the mature neuronal marker NeuN (green). Nuclei were counterstained with DAPI (blue). Scale bars, 100 μm. **(D-E)** Quantification of neuronal differentiation at day 21. **(D)** βIII-tubulin$^+$ area fraction (%). **(E)** Percentage of TBR1$^+$ cells (TBR1$^+$/DAPI$^+$). **(F)** Representative immunofluorescence images of day 28 hCOs labeled with EdU (green) and DAPI (blue) to assess EdU incorporation. Scale bars, 100 μm. **(G)** Quantification of EdU$^+$ cells at day 28 as the percentage of EdU$^+$ cells (EdU$^+$/DAPI$^+$). Data are presented as mean ± SEM. Each dot represents one organoid. Statistical significance was determined using a two-tailed Student's t-test. *P < 0.05, **P < 0.01, ***P < 0.001.

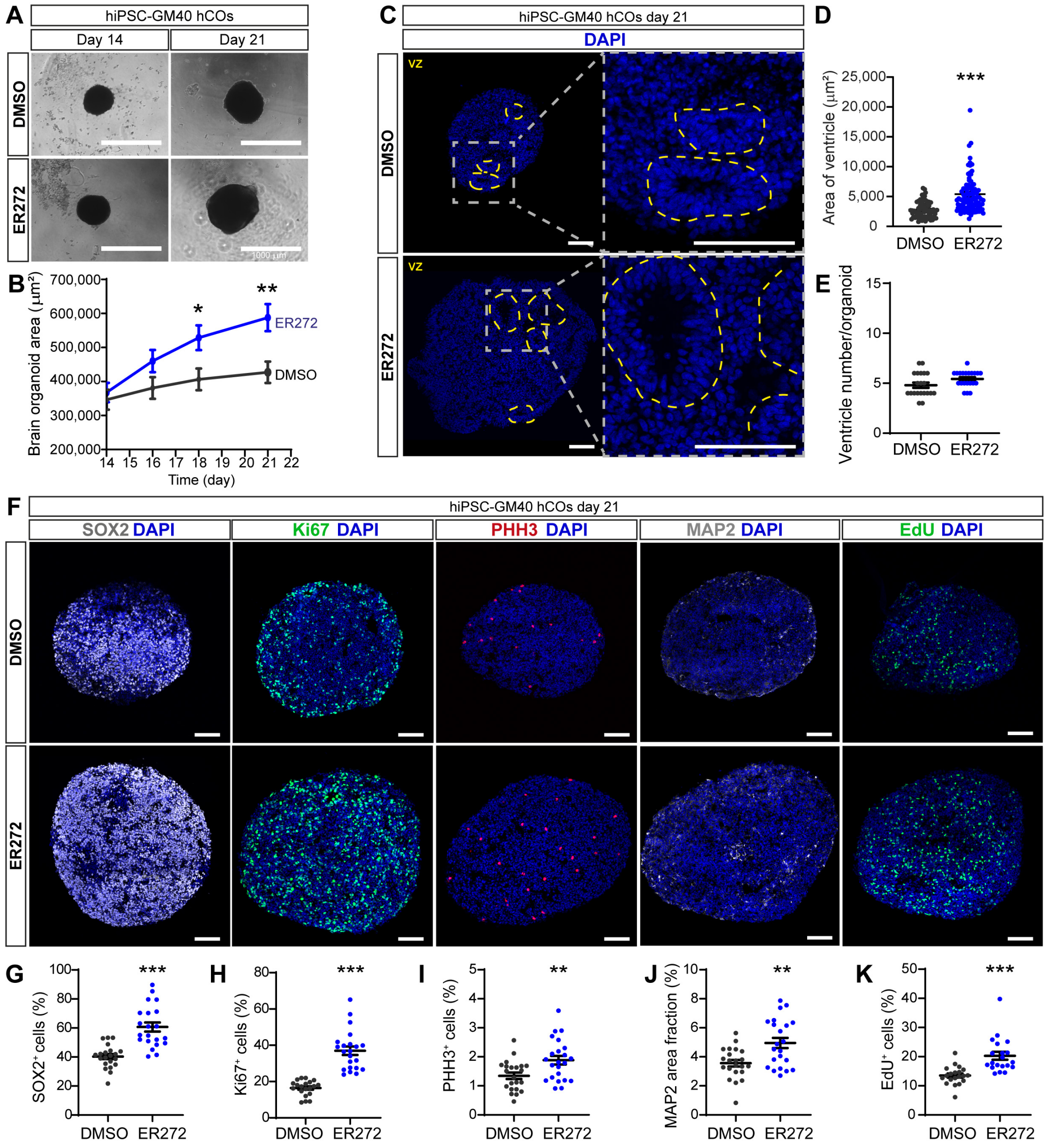


**Supplementary Figure 2. ER272 promotes ventricular zone expansion, neural progenitor proliferation, and neuronal differentiation in hiPSC-derived human cortical organoids (hCOs). (A)** Representative bright-field images of GM40 hiPSC-derived human cortical organoids (hiPSC-GM40 hCOs) cultured in the presence of vehicle (DMSO) or ER272 from day 14 to day 21. Scale bars, 1000 µm. **(B)** Quantification of organoid area during the treatment period (days 14-21), showing enhanced organoid growth following ER272 treatment. **(C)** Representative DAPI-stained sections of day 21 hiPSC-GM40 hCOs illustrating ventricular zone (VZ)-like structures (outlined by dashed yellow lines). Higher-magnification images show representative VZ-like regions. Scale bars, 100 µm. **(D-E)** Quantification of ventricular zone-like structures at day 21. **(D)** Area of ventricular zone-like structures (µm$^2$). **(E)** Number of ventricular zone-like structures per organoid. **(F)** Representative immunofluorescence images of day 21 hiPSC-GM40 hCOs stained for the neural progenitor marker SOX2 (gray), the proliferation marker Ki67 (green), the mitotic marker phospho-histone H3 (PHH3; red), the neuronal marker MAP2 (gray), and EdU (green). Nuclei were counterstained with DAPI (blue). Scale bars, 100 µm. **(G-K)** Quantification of neural progenitor, proliferation, and neuronal differentiation markers. **(G)** Percentage of SOX2$^+$ neural progenitors (SOX2$^+$/DAPI$^+$). **(H)** Percentage of proliferating Ki67$^+$ cells (Ki67$^+$/DAPI$^+$). **(I)** Percentage of mitotic PHH3$^+$ cells (PHH3$^+$/DAPI$^+$). **(J)** Neuronal MAP2$^+$ area fraction (%). **(K)** Percentage of EdU$^+$ cells (EdU$^+$/DAPI$^+$). Data are presented as mean ± SEM. Each dot represents one organoid. Statistical significance was determined using a two-tailed Student's t-test. *P < 0.05, **P < 0.01, ***P < 0.001.

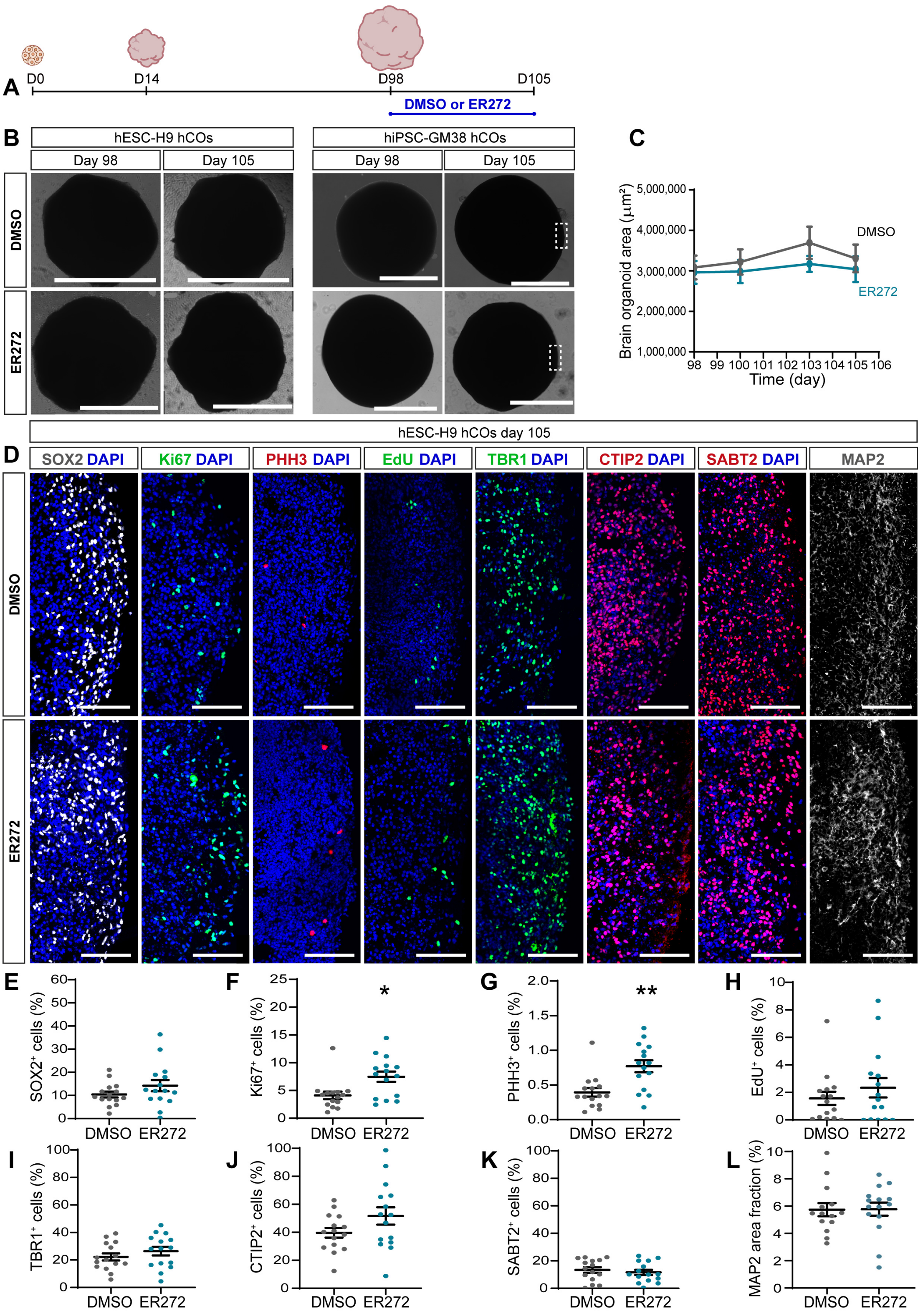


**Supplementary Figure 3. Characterization of 3.5-month-old human cortical organoids (hCOs) following ER272 treatment. (A)** Schematic representation of the experimental timeline for H9 hESC and GM38 hiPSC-derived late-stage hCOs. hCOs were cultured until day 98 (D98) and subsequently treated with DMSO (vehicle) or ER272 for 7 days before analysis at day 105 (D105). **(B)** Representative bright-field images of hiPSC-GM38 and hESC-H9 hCOs treated with DMSO or ER272 from day 98 to day 105. Dashed boxes indicate representative regions shown at higher magnification in (D). **(C)** Quantification of organoid area over time in 3.5-month-old hCOs showing no significant effect of ER272 treatment on organoid growth. **(D)** Immunofluorescence images of day-105 hESC-H9 hCOs stained for SOX2, KI67, PHH3, EdU, TBR1, CTIP2, SATB2, and MAP2. Nuclei were counterstained with DAPI. **(E-L)** Quantification of SOX2⁺, KI67⁺, PHH3⁺, EdU⁺, TBR1⁺, CTIP2⁺, SATB2⁺, and MAP2⁺ area fraction. Data are presented as mean ± SEM. *$p < 0.05$, **$p < 0.01$. Scale bars, 1000 µm (low magnification) and 100 µm (high magnification).

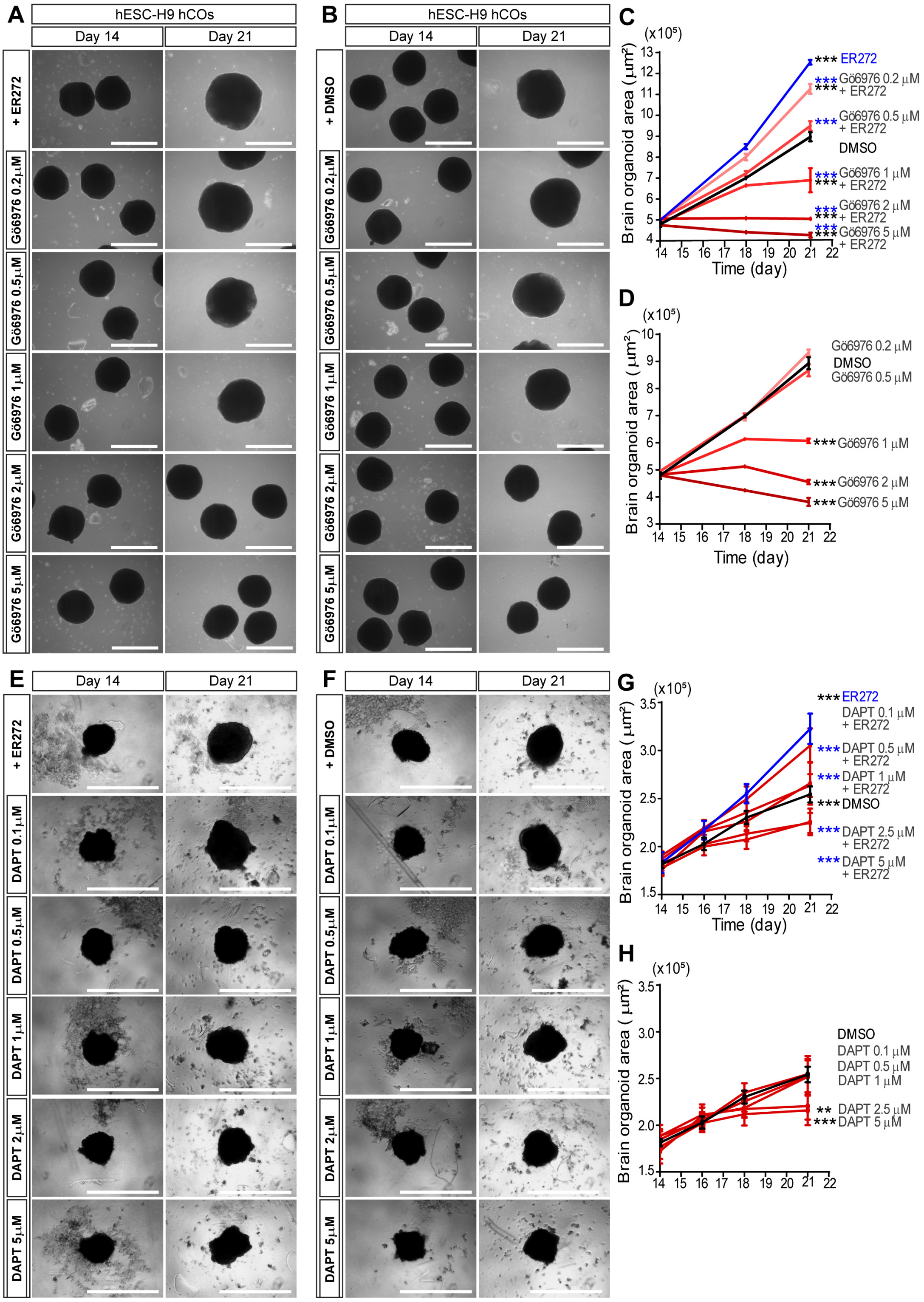


**Supplementary Figure 4. PKC and Notch pathways play an important role in controlling basal human brain organoid (hCOs) growth. (A-B)** Bright-field images of H9 human embryonic stem cell-derived cortical organoids (hESC-H9 hCOs) cultured from day 14 to day 21 in the presence of vehicle (DMSO) or ER272 combined with increasing concentrations of the classical PKC inhibitor Gö6976. Scale bar, 1000 μm. **(C)** Quantification of organoid area during treatment with DMSO or Gö6976 alone, showing a dose-dependent reduction in organoid growth following PKC inhibition. **(D)** Quantification of organoid area during treatment with ER272 in combination with increasing concentrations of Gö6976. PKC inhibition progressively attenuated the growth-promoting effect of ER272. **(E-F)** Bright-field images of hESC-H9 hCOs cultured from day 14 to day 21 in the presence of vehicle (DMSO) or ER272 combined with increasing concentrations of the γ-secretase/Notch inhibitor DAPT. **(G)** Quantification of organoid area during treatment with DMSO or increasing concentrations of DAPT, showing the effect of Notch pathway inhibition on organoid growth. **(H)** Quantification of organoid area during treatment with ER272 in combination with increasing concentrations of DAPT. Inhibition of Notch signaling progressively reduced the growth-promoting effect of ER272. Data are presented as mean ± SEM. Statistical significance was determined using two-way ANOVA. *$P < 0.05$, **$P < 0.01$, ***$P < 0.001$.

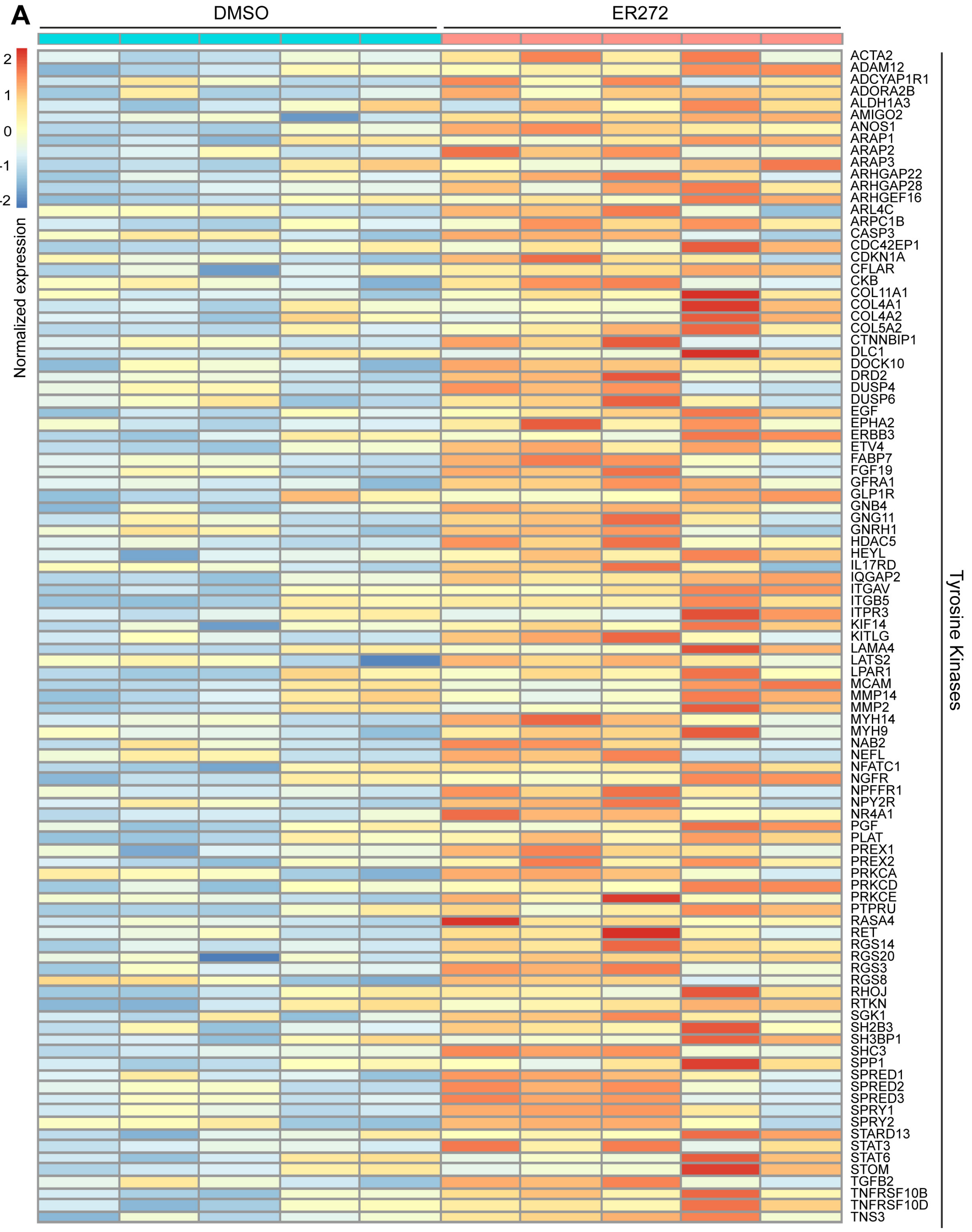


**Supplementary Figure 5. ER272 upregulates genes associated with receptor tyrosine kinase signaling in human cortical organoids. (A)** Heatmaps of the upregulated genes associated with tyrosine kinases pathway in ER272-treated hCOs compared with DMSO-treated hCOs. VST-transformed read counts are shown, with each row representing one gene and each column one independent hCO sample.

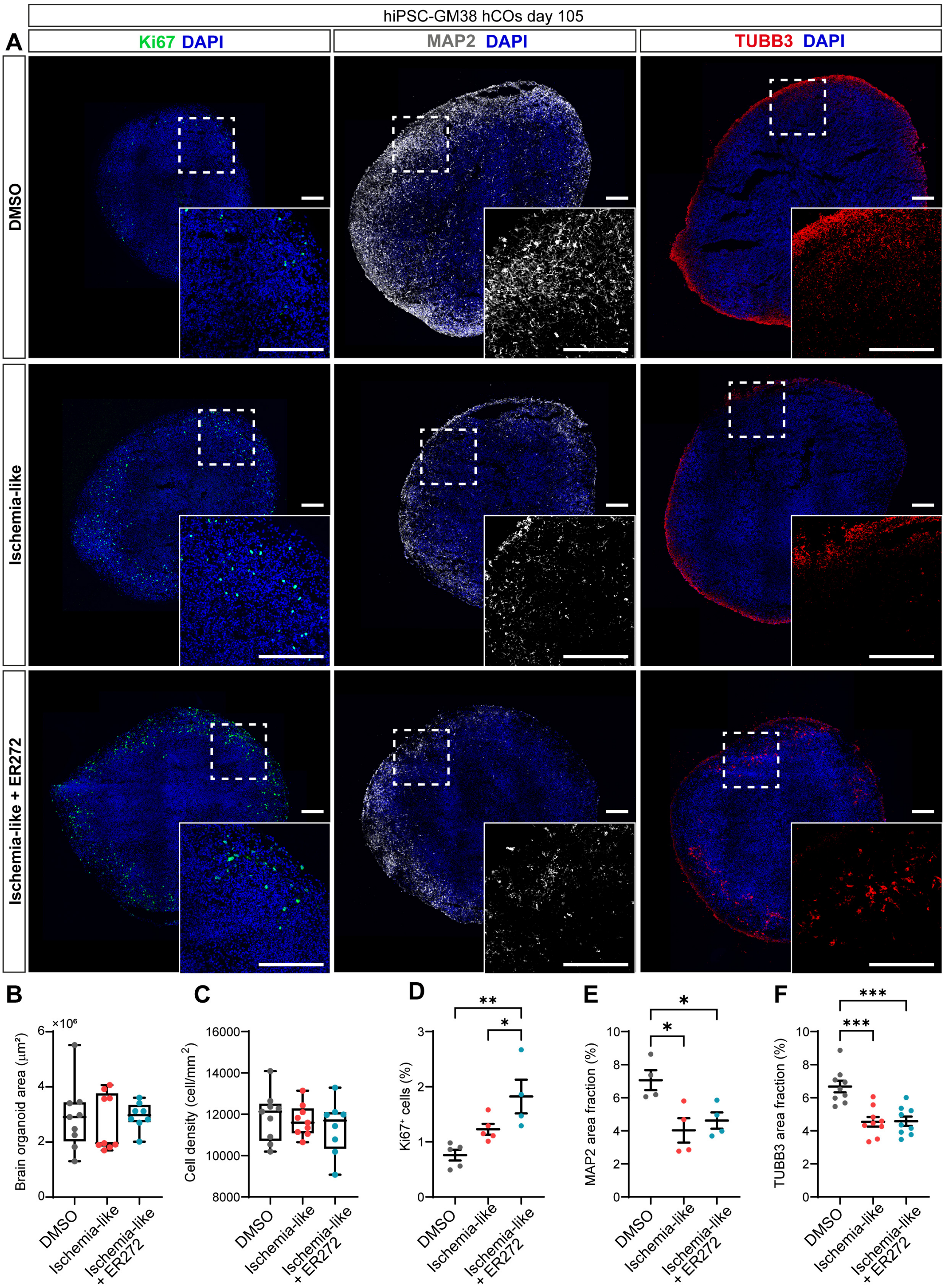


**Supplementary Figure 6. Effects of ER272 treatment on human cortical organoids following an ischemia-like insult.** **(A)** Representative immunofluorescence images of day-105 hiPSC-GM38 hCOs treated with DMSO, subjected to an ischemia-like injury, or subjected to an ischemia-like injury followed by ER272 treatment. hCOs were stained for Ki67, MAP2, and β-III-Tubulin. Nuclei were counterstained with DAPI. Dashed boxes indicate representative regions shown at higher magnification in the corresponding insets. Scale bars, 200 µm. **(B-C)** Quantification of organoid area and cell density, respectively. **(D-F)** Quantification of proliferating Ki67/DAPI-positive cells, neuronal MAP2 area fraction, and neuronal β-III-Tubulin area fraction, respectively. Data are presented as mean ± SEM. *P < 0.05, **P < 0.01, ***P < 0.001.

**SUPPLEMENTAL TABLES**

**Supplementary Table 1. List of differentially expressed genes (DEGs) in DMSO-treated versus ER272-treated hCOs at day 21.** The separate worksheets contain: (i) all DEGs with an adjusted p value (padj) < 0.05, (ii) upregulated DEGs, and (iii) downregulated DEGs. For each gene, the table includes ENSEMBL gene ID, mean normalized expression (baseMean), log2 fold change (log2FoldChange), standard error of log2 fold change (lfcSE), p-value, adjusted p-value (padj), and gene symbol (GeneName).

**Supplementary Table 2. Lists of enriched pathways using the Reactome Pathways 2024 database in DEGs from DMSO-treated versus ER272 hCOs at day 21.** Separate worksheets include enrichments for upregulated DEGs.

### MATERIALS AND METHODS

#### Culture of human embryonic stem cells (hESCs) and induced pluripotent stem cells (hiPSCs)

Human embryonic stem cells (hESCs; H9/WA09, WiCell; metadata: female, no disease associated) and human induced pluripotent stem cells (hiPSCs; GM23338 and GM23340*A, Coriell; metadata: males, no disease associated) were maintained under feeder-free conditions. Cells were cultured in mTeSR Plus medium (StemCell Technologies 100-0276) on Geltrex-coated plates (ThermoFisher A1413302) and routinely passaged using EDTA (Sigma 03690). Following thawing, hESC and hiPSC cultures were maintained for no more than 10-15 additional passages. hESC and hiPSC lines were authenticated by WiCell upon acquisition, and pluripotency was routinely assessed by qPCR and immunofluorescence analysis of pluripotency markers. Mycoplasma testing was performed every 3 months. All cultures were maintained without antibiotics under good laboratory practices. Genomic characterization was performed by copy-number variation analysis (qPCR CNV) using genomic DNA extracted from cells derived from cryopreserved stocks at approximately passages 30-40.

#### Human cortical organoid (hCO) differentiation

Cortical brain organoids were developed following a modified version of the Sasai protocol [33]. hESC-H9 and hiPSC-GM38/GM40 cell lines were dissociated using accutase (Stem Cell Technologies 7922) to obtain a suspension of individual cells that were seeded into 96 well U bottom plates (Nunclon sphera 15396123) at 9,000 cells per well. Plates were centrifuged 2 min at 550 rpm to cluster the cells at the bottom of the well. Organoids were cultured in DMEM-F12 (ThermoFisher 31331-093) with 20% KO serum (Fisher 10828028), supplemented with 1/100 non-essential amino acid (NEAA; ThermoFisher 11140-035), 1/100 penicillin/streptavidin (Pen/strep; ThermoFisher 15140122) and 100µM β-mercaptoethanol (Sigma 444203) for 14 days. Morphogens were added to the medium for 14 days: 5µM SB (SB431542; Sigma S4317) and 3µM IWR1 (Sigma ref: 681669). Rock inhibitor (Y-27632 2HC; Bio-Connect S1049) was added on day 0 when seeding the cells at 20µM and then at 10µM from day 2 to day 6. From day 14 to day 28, medium was composed of DMEM-F12, 1/100 sodium pyruvate (Na Pyr; ThermoFisher 11360-039), 1/100 NEAA, 1/100 N2 (ThermoFisher 17502048), 1/100 Pen/strep, 1/150 BSA (BSA fraction V; ThermoFisher 9048-46-8), 1/50 B27 without vitamin A (ThermoFisher 11500446) and 100µM β-mercaptoethanol. Organoids were transferred to 6cm dishes (Greiner CLS430196) when their area was around 0,7mm$^2$ and placed on orbital shaker (75 rpm). From day 28 to day 35, medium was composed of half DMEM-F12 and half Neurobasal (ThermoFisher 21103-049) and supplemented with 1/200 Na Pyr, 1/200 NEAA, 1/200 N2, 1/200 Glutamax (ThermoFisher 35050-038), 1/300 BSA, 1/100 Pen/strep, 1/50 B27 without vitamin A and 50µM β-mercaptoethanol. From day 35 till the end of the culture, the medium was similar to day 28 to day 35 but supplemented with B27 with vitamin A (ThermoFisher 11530536). Organoids were transferred to bioreactor tubes (Bioké 2800005) when their area was around 1.4 mm$^2$ and kept in culture in a bioreactor at 75 rpm (Bioké 2800000). At day 70, 1% of Matrigel (Corning 354230) was added to the culture.

Medium was changed every 2-3 days and organoid pictures were acquired on the same days to assess area and overall quality till their transfer to the bioreactors.

**Isolation of ER272**

The isolation and purification of 12-deoxyphorbol 13-isobutyrate, also known as DPB [43] or ER272 [14,15] (CAS: 25090-74-8) were carried out in our laboratory according to the procedures described in the supplementary methods of Geribaldi-Doldan et al. [13].

**Reagents and hCO drug treatments in culture**

Human cortical organoids derived from hESC-H9 or from hiPSC-GM40/GM38 were treated with either DMSO (vehicle) or 1μM ER272 during 7 days from day 14 to day 21 or from day 98 to day 105. Additionally, DMSO or ER272 treated hESC-H9 hCOs were cultured in the presence of classical PKC inhibitor Gö6976 (Sigma-Aldrich 365250) or the γ-secretase inhibitor DAPT (Sigma D5942) to prevent downstream Notch signaling from day 14 to day 21. In addition, hCOs were cultured in the presence of 5-ethynyl-2'-deoxyuridine (1μM EdU, Invitrogen C-10337) as indicated in the corresponding figure.

**Oxygen and glucose deprivation (OGD)**

To mimic an ischemia-like brain injury, day-98 hCOs, were deprived of glucose and oxygen for 6 h. Oxygen and glucose deprivation (OGD) was achieved by incubating hCOs in glucose-free medium (DMEM/F12, without L-glutamine, without HEPES, without glucose, VWR) in a humidified anaerobic chamber Panasonic Healthcare containing 5% $CO_2$, 94.9% N2 and 0.1% $O_2$ at 37°C for 6 h. After OGD exposure, hCOs were returned to regular hCO culture medium under oxygen conditions. Cultures were maintained for an additional week. ER272 (1μM) or vehicle was administered during this period according to the experimental condition. Control hCOs were subjected to the same manipulations but kept under normoxia and regular hCO culture medium containing glucose.

**Organoid-to-organoid transplantation model**

An organoid-to-organoid transplantation model was established by gently positioning 2-week-old GFP-labeled early cortical organoids (hCOs) in contact with the surface of 98-day-old late-stage hCOs for 7 days or 14 days *in vitro*. Organoid-to-organoid transplantation was evaluated under three conditions: (i) control treated with DMSO, (ii) OGD-exposed treated with DMSO, and (iii) OGD-exposed treated with ER272 (1μM). After transplantation, brightfield and fluorescence live images of the GFP-grafted organoids were acquired every 2-3 days for all experimental conditions. Imaging was performed using a Carl Zeiss inverted microscope equipped with a digital camera, using ZEN Lite software. Images were captured using a 5× objective. After collection and fixation, total graft area ($\mu m^2$) and the extent of graft integration into the host organoid was evaluated. Graft size was defined as the total transplanted GFP-organoid area ($\mu m^2$), whereas graft integration was quantified as the GFP-positive area integrated into the host organoid area.

**Organoid processing and immunostaining**

Organoids were rinsed in phosphate-buffered saline solution (PBS), fixed in 4% paraformaldehyde solution in PBS (PFA) for 45 min at 4°C, then transferred to 30% sucrose solution in PBS for 48 h at 4°C. Excess sucrose solution was removed from organoids by moving them through embedding medium (Epredia, Richard-Allan Scientific, MI, USA) after which they were flash frozen and stored at -80°C until further use. Blocks were cut on a Cryostar NX70 cryostat into 14 µm thick sections collected onto slides (Superfrost Plus Adhesion, Epredia J1800AMNZ), 3 to 4 series per slide, air-dried overnight, and kept at -20°C until staining.

Immunofluorescence of organoid cryosections was performed directly on the slides as described before [39]. Slides were first rinsed in PBS for 5 min to remove embedding medium, followed by antigen retrieval in target retrieval solution (Dako S1699) at 90°C for 15 min. Slides were then rinsed 3 x 5 min in a bath of PBS at room temperature. Permeabilization was done in 5% BSA (Carl Roth 0163.2), 5% donkey serum, and 0.3% Triton X-100 in PBS for 30 min at room temperature by adding 500 µL of solution to each slide. Afterwards, primary antibodies were added in a buffer solution of 3% BSA, 1% donkey serum, and 0.1% Triton, in PBS, at 350 µL per slide and slides were incubated at 4°C overnight in a wet box and covered with Parafilm. The next day, slides were washed 3 times in PBS and incubated for 3 h in a dark box at RT with secondary antibodies in buffer solution at 1/500. Slides were then washed 2 times 5 min with 500 µL PBST (0.1% Tween-20 (Sigma 9036-19-5) in PBS) and 1 time with PBS and then incubated in DAPI solution (2 ng/ml) for 20 min. 5-ethynyl-2deoxyuridine (EdU) labeled DNA was detected using Click-iT EdU cell proliferation kit for Imaging, Alexa Fluor 488 dye following the manufacturer's protocol (Invitrogen C-10337). Slides were washed 2 times in PBS, cover-slipped using fluorescent mounting agent (Dako S3023, Agilent Technologies Inc, Santa Clara, CA, USA), and air-dried for 48h. Immunostained slides were stored at 4°C for imaging and -20°C for long-term storage. All primary and secondary antibodies used are listed in the tables below.

| Primary Abs | Host | Company | Reference | Dilution |
|---|---|---|---|---|
| β-Actin HRP | Mouse | Cell signaling | 12262S | 1/10000 |
| β-III-Tubulin | Guinea pig | Synaptic Systems | 302 304 | 1/500 |
| Cleaved Caspase-3 | Rabbit | Cell signaling | 9661 | 1/400 |
| CTIP2 | Rat | Abcam | ab18645 | 1/300 |
| GFAP | Rabbit | Agilent Dako | Z033429 | 1/500 |
| GFP | Goat | Abcam | ab6673 | 1/500 |
| HuCD | Mouse | Invitrogen | A-21271 | 5 ug/mL |
| Ki67 | Rabbit | Abcam | ab15580 | 1/500 |
| MAP2 | Chicken | Abcam | ab5392 | 1/2000 |
| Nestin | Mouse | Abcam | ab22035 | 1/1000 |
| NeuN | Rabbit | Cell Signaling | 12943 | 1/500 |
| NICD | Rabbit | Cell Signaling | 4147 | 1/500 |
| PHH3 | Mouse | Invitrogen | MA5-15220 | 1/300 |
| SATB2 | Mouse | Abcam | ab51502 | 1/500 |
| SOX2 | Mouse | BD Biosciences | 561469 | 1/100 |
| TBR1 | Rabbit | Abcam | ab31940 | 1/250 |

| Secondary Abs | Host | Company | Reference | Dilution |
|---|---|---|---|---|
| AF488 anti-goat | Donkey | Invitrogen | a-11055 | 1/500 |
| AF488 anti-rabbit | Donkey | Invitrogen | a-21206 | 1/500 |
| AF488 anti-rat | Donkey | Invitrogen | a-21208 | 1/500 |
| AF555 anti-guinea pig | Goat | Invitrogen | a-21435 | 1/500 |
| AF555 anti-mouse | Donkey | Invitrogen | A32773 | 1/500 |
| AF555 anti-rabbit | Donkey | Invitrogen | a-31572 | 1/500 |
| AF555 anti-rat | Donkey | Invitrogen | a-78945 | 1/500 |
| AF647 anti-chicken | Donkey | Invitrogen | a-78952 | 1/500 |
| AF647 anti-mouse | Donkey | Invitrogen | a-31571 | 1/500 |
| AF647 anti-rabbit | Donkey | Invitrogen | a-31573 | 1/500 |
| HRP anti-rabbit | Goat | Cell signaling | 7074 | 1/2000 |
| HRP anti-mouse | Goat | Cell signaling | 7076 | 1/2000 |
| HRP anti-chicken | Goat | Invitrogen | a-16054 | 1/2000 |

**Imaging and image analysis**

Immunostained organoid sections were imaged with a 20x, plan-apochromat objective on a Nikon A1R confocal microscope. XY scanning stage was used for tile scans with 405 nm, 488 nm, 555 nm, and 647 nm lasers. The NIS-Element AR software was utilized to automatically stitch together individual images with a 15% overlap of the field of view. Analysis was performed using the Fiji package of ImageJ software (U.S. National Institutes of Health).

**Quantification of transplant integration**

Transplant integration was quantified based on the GFP$^+$ area of the transplanted organoid using ImageJ/Fiji. First, the entire GFP$^+$ transplant area was delineated and measured, corresponding to the total transplant area. In parallel, the area of the GFP$^+$ transplant located within the host organoid was also delineated and measured, corresponding to the integrated transplant area. The percentage of transplant integration was calculated as the area of the GFP$^+$ transplant located within the host organoid divided by the total GFP$^+$ transplant area, expressed as a percentage.

**Quantification of % SOX2, Ki67, PHH3, EdU, TBR1, CTIP2, SATB2, NeuN, cleaved Casp3, HuCD**

Quantification of SOX2$^+$, Ki67$^+$, PHH3$^+$, EdU$^+$, TBR1$^+$, and CTIP2$^+$ cells was performed semi-automatically using ImageJ/Fiji. First, background fluorescence was removed using the Subtract Background function. Binary masks were then generated from each fluorescence channel using marker-specific thresholding methods (Moments for SOX2, Ki67, PHH3, CTIP2, SATB2 and NeuN; Default for EdU and DAPI; Li for CTIP2, and manual thresholding for HuCD). Threshold values were selected by visual comparison of the resulting binary image with the original 16-bit image to ensure accurate representation of the staining signal. Positive cells were identified using the Analyze Particles function with a minimum particle size of 10 $\mu m^2$. The number of positive cells was expressed as a percentage of the total number of DAPI$^+$ nuclei.

For CTIP2$^+$EdU$^+$ quantification, double-positive cells were identified manually and normalized to the total number of EdU$^+$ nuclei.
For cleaved Casp3 quantification, positive cells were identified manually and expressed as a percentage of the total number of DAPI$^+$ nuclei.

**Quantification of % organoid area stained for β-III-tub, MAP2, GFAP**
Quantification of β-III-tubulin, MAP2, and GFAP markers was performed automatically using a custom ImageJ/Fiji macro. First, the total organoid area was segmented from the DAPI channel using the Huang thresholding method, allowing the entire organoid to be identified as a single object. The resulting organoid mask was then applied to the channel corresponding to the marker of interest, restricting the analysis to the organoid area. Positive regions were segmented using the Moments thresholding method. Finally, the percentage of marker-positive area relative to the total organoid area was quantified using the ImageJ "% Area Fraction" measurement, which calculates the percentage of the total organoid area occupied by the positive signal.

**Quantification of number of GFP$^+$ cells and fibers**
The number of GFP$^+$ cells and fibers was quantified using ImageJ/Fiji. The host organoid area was first selected to define the region of interest. Background fluorescence was then removed using the Subtract Background function, followed by thresholding using the Default method. The resulting binary mask was used to quantify GFP$^+$ cells and fibers within the selected host area. A minimum particle size of 15 $\mu m^2$ was applied during particle analysis. The number of GFP$^+$ particles was then normalized to the area of the corresponding host organoid to account for differences in host size and expressed as GFP$^+$ particles per area.

**Protein extracts and Immunoblotting**
Brain organoids were homogenized and lysed on ice using in RIPA buffer (50mM Tris-HCl, 150mM NaCl, 1mM EDTA, 1% Triton X-100, 1% Na Deoxycholate, 0.1% SDS) and Complete Protease Inhibitor Cocktail (Merck Millipore 11697498001) and phosphatase inhibitor cocktail (Merck 11697498001). Proteins concentration was assessed using a Pierce BCA protein Assay Kit (ThermoFisher 23227). BCA standards have been prepared according to the manufacturer and samples have been diluted 1/10 or 1/4. Ten µl of standards or samples have been mix with 200µl of working solution prepared according to the manufacturer and incubated for 30 min at 37°C. Absorbance have been measure using a spectrophotometer (562 nm). Concentrations of the samples were determined manually based on the equation from the standards curve. Samples (30µg) were mixed with Sample Buffer, Laemmli 2× Concentrate (Sigma S3401) and heated at 70°C for 10 min. Samples and ladder (ThermoFisher 26619) were loaded on bis-tris 4-12% precast gels (ThermoFisher NP0322BOX) in MOPS SDS running buffer 1X (ThermoFisher NP0001) for about 1h30 at 120V. Proteins were transferred on a PVDF membrane (Merck GE10600023) on ice cold transfer buffer (25mM Tris, 192 mM glycine and 20% (v/v) methanol) for 1h15 at 300mA. A Ponceau staining was done to ensure appropriate transfer and the membrane washed 3 times 5 min in TBTS 1X (150mM NaCl, 50mM Tris base, 0,1% Tween, pH =7,5) to remove the coloration. Membrane was blocked in 5% milk in TBST 1X for 1h and incubated with primary antibody (1/1000) in 1% milk in TBST 1X overnight. Three washes of 10 min were

performed to remove excess of primary antibody and secondary antibody rabbit, mouse, or chicken HRP (Cell signaling 7074, 7076; and Invitrogen A16054 respectively) was added at 1/1000 in 1% milk in TBST 1X and incubated for 1 h. After 3 washes in TBST 1X, the membrane was revealed using Pierce ECL WB substrate (ThermoFisher 32106). After the picture for the protein of interest, the membrane was washed 3 times in TBST 1X and incubated for 1 h in β-actin HRP (dilution 1/10000, Cell Signaling 12262S). The membrane was revealed a second time to take the picture of the β-actin.

**RNA isolation and sequencing, data analysis and differential expression analysis**
After treatment, early-stage hCOs were collected at day 21. RNA was extracted using the RNeasy mini kit (Qiagen 74104), followed by a DNase treatment to remove genomic DNA (Ambion AM2222). Sample quality was assessed using a Nanodrop 2000 (ThermoFisher). High-quality RNA samples with RNA integrity number (RIN) values of 8-10 were used for sequencing. hESC-H9 hCOs treated with DMSO (n=5) or treated with ER272 (n=5) from 2 independent differentiations were analyzed. The total extracted RNA was analyzed by the GIGA Genomic platform (https://www.gigagenomics.uliege.be) and generated the polyA RNA-seq libraries using Illumina Stranded Total RNA Prep Ligation with Ribo-Zero Plus following the manufacturer protocol (Illumina 20040529). The libraries were dosed through qPCR using the KAPA SYBR Fast* Mastermix (2X) kit Universal (Sopachem KK4602) and were sequenced using the Illumina Novaseq 6000 sequencer with the flowcell Novaseq S4 V1.5 (300 cycles), generating 2 x 150-bp paired-end reads. Raw reads were processed through nf-core/RNAseq pipeline (version 3.0) using human genome build GRCh38 and the annotations from Ensembl release 103 (ensembl.org). Using the raw counts as input, R package DESeq2 (v1.20.0) was used to perform the normalization, clustering and differential expression analysis and obtain the differentially expressed genes (DEGs). RUVSeq (v1.20.0) was used to remove unwanted variation (batch correction). Enrichment analysis for DEGs was conducted with EnrichR.

**Statistical analysis**
All statistical analysis was performed using GraphPad (Prism). For each experiment, the definition of the experimental unit, the number of organoids analyzed and the number of independent experiments are indicated in the corresponding figure legend. Data were first tested for normality with a Shapiro-Wil test. For comparisons between two groups, an unpaired, two-tailed Student's *t*-test was used for normally distributed data, whereas the corresponding non-parametric Mann–Whitney *U* test was used when the data did not follow a normal distribution. For comparisons involving more than two groups, one-way ANOVA followed by Tukey's multiple-comparisons test was used for normally distributed data, whereas the Kruskal-Wallis test followed by Dunn's multiple-comparisons test was used for non-normally distributed data. For experiments involving two independent variables, two-way ANOVA followed by Šídák's multiple-comparisons test was used. The specific statistical test, and sample size ($n$) for each analysis are indicated in the corresponding figure legends. Statistical significance was defined as $p < 0.05$; p < 0.01; $p < 0.001$.
